# Convergent network effects along the axis of gene expression during prostate cancer progression

**DOI:** 10.1101/2020.02.16.950378

**Authors:** Konstantina Charmpi, Tiannan Guo, Qing Zhong, Ulrich Wagner, Rui Sun, Nora C. Toussaint, Christine E. Fritz, Chunhui Yuan, Hao Chen, Niels J. Rupp, Ailsa Christiansen, Dorothea Rutishauser, Jan H. Rüschoff, Christian Fankhauser, Karim Saba, Cedric Poyet, Thomas Hermanns, Kathrin Oehl, Ariane L. Moore, Christian Beisel, Laurence Calzone, Loredana Martignetti, Qiushi Zhang, Yi Zhu, María Rodríguez Martínez, Matteo Manica, Michael C. Haffner, Ruedi Aebersold, Peter J. Wild, Andreas Beyer

## Abstract

**Background:** Tumor-specific genomic aberrations are routinely determined by high throughput genomic measurements. It remains unclear though, how complex genome alterations affect molecular networks through changing protein levels, and consequently biochemical states of tumor tissues.

**Results:** Here, we investigated the propagation of genomic effects along the axis of gene expression during prostate cancer progression. For that, we quantified genomic, transcriptomic and proteomic alterations based on 105 prostate samples, consisting of benign prostatic hyperplasia regions and malignant tumors, from 39 prostate cancer patients. Our analysis revealed convergent effects of distinct copy number alterations impacting on common downstream proteins, which are important for establishing the tumor phenotype. We devised a network-based approach that integrates perturbations across different molecular layers, which identified a sub-network consisting of nine genes whose joint activity positively correlated with increasingly aggressive tumor phenotypes and was predictive of recurrence-free survival. Further, our data revealed a wide spectrum of intra-patient network effects, ranging from similar to very distinct alterations on different molecular layers.

**Conclusions:** This study uncovered molecular networks with remarkably convergent alterations across tumor sites and patients, but it also exposed a diversity of network effects: we could not identify a single sub-network that was perturbed in all high-grade tumor regions.

## Background

Prostate cancer (PCa) represents one of the most common neoplasms among men with almost 1,300,000 new cases and 360,000 deaths in 2018 ^1^ accounting for 15% of all cancers diagnosed. PCa is the fifth leading cause of cancer death in men and represents 6.6% of total cancer mortality in men [1]. Despite earlier detection and new treatments, the lifetime risk to die of PCa has remained stable at approximately 3% since 1980. (National Cancer Institute SEER data: https://seer.cancer.gov/statfacts/html/prost.html). In many patients, PCa is indolent and slowly growing. The challenge is to identify those patients who are unlikely to experience significant progression while offering radical therapy to those who are at risk. Current risk stratification models are based on clinicopathological variables including histomorphologically defined grade groups, prostate-specific antigen (PSA) levels and clinical stage. Although those variables provide important information for clinical risk assessment and treatment planning [2, 3], they do not sufficiently predict the course of the disease.

Extensive genomic profiling efforts have provided important insights into the common genomic alterations in primary and metastatic PCa [4-9]. Interestingly, PCa genomes show a high frequency of recurrent large-scale chromosomal rearrangements such as TMPRSS2-ERG [10]. In addition, extensive copy number alterations (CNAs) are common in PCa, yet point mutations are relatively infrequent in primary PCa compared to other cancers [6, 11]. A major complicating factor is that around 80% of PCas are multifocal and harbor multiple spatially and often morphologically distinct tumor foci [12, 13]. Several recent studies have suggested that the majority of topographically distinct tumor foci appear to arise independently and show few or no overlap in driver gene alterations [14-16]. Therefore, a given prostate gland can harbor clonally independent PCas.

To allow for a more functional assessment of the biochemical state of PCa, it is necessary to go beyond genomic alterations and comprehensively catalogue cancer specific genomic, transcriptomic and proteomic alterations in an integrated manner [17-19]. Such an approach will provide critical information for basic and translational research and could result into clinically relevant markers. While hundreds of PCa genomes and transcriptomes have been profiled to date [20], little is known about the PCa proteome. Although recent work has emphasized the need for integrated multi-omics profiling of PCa, we still lack understanding about how genomic changes impact on mRNA and protein levels [17-19]. Especially the complex relationship between tumor grade, tumor progression and multi-layered molecular network changes remains largely elusive.

For example, previous work has shown that copy number changes may alter transcript levels of many genes, whereas the respective protein levels remain relatively stable [21]. Indeed, there is compelling evidence across multiple tumor types that many genomic alterations are ‘buffered’ at the protein level and are hence mostly clinically inconsequential [22]. To better understand the evolution of PCa and to identify core networks perturbed by genomic alterations and thus central for the tumor phenotype, it is therefore essential to investigate the transmission of CNAs to the transcriptomic and proteomic level.

To this end, it is important to decipher which genomic alterations impact PCa proteomes, which of those proteomic alterations are functionally relevant, and how molecular networks are perturbed at the protein level across tumors.

To address these open questions, we performed a multi-omics profiling of radical prostatectomy (RP) specimens at the level of the genome, transcriptome and proteome from adjacent biopsy-level samples, using state-of-the-art technologies. Unique features of this study are (1) the utilization of PCT (pressure cycling technology)-SWATH (Sequential Window Acquisition of all THeoretical Mass Spectra) mass spectrometry [23, 24], allowing rapid and reproducible quantification of thousands of proteins from biopsy-level tissue samples collected in clinical cohorts; (2) the simultaneous profiling of all omics layers from the same tissue regions; (3) inclusion and full profiling of benign regions, which provides a matching control for each tumor; and (4) the full multi-omics characterization of multiple tumor regions from the same patients, thus enabling the detailed investigation of tumor heterogeneity. This design resulted in the multi-layered analyses of 105 samples from 39 PCa patients, as well as of the exome of corresponding peripheral blood cells yielding a comprehensive molecular profile for each patient and identified molecular networks that are commonly altered in multiple patients. Importantly, some of the affected genes/proteins exhibited very small individual effect sizes, suggesting that combined network effects of multiple genes may significantly contribute to determining PCa phenotypes.

## Results

### Proteogenomic analysis of the sample cohort identifies known PCa biomarkers

In this study, we analyzed 39 PCa patients (**Additional file 1: Fig. S1**) belonging to three groups who underwent laparoscopic robotic-assisted RP. The patients were from the PCa Outcomes Cohort (ProCOC) study [25, 26]. Tumor areas were graded using the ISUP (International Society of Urological Pathology) grade groups [27], which range from ISUP grade group G1 (least aggressive) to G5 (most aggressive). The more advanced grade groups G4 and G5 were considered jointly (G4/5). The cohort tested included 12 low-grade (G1), 17 intermediate-(G2 and G3), and 10 high-grade (G4/5) patients (**Fig. 1a, Additional file 1: Fig. S1, Additional file 2: Table S1**). For low-grade PCa patients, we selected two representative regions, one of benign prostatic hyperplasia (BPH) and one of malignant tumor (TA). Since PCa often presents as a multifocal disease with heterogeneous grading within each prostate specimen [24] we analyzed two different tumor regions from the 27 intermediate-and high-grade patients. In those cases three representative regions, including BPH, the most aggressive tumor (TA1) and a secondary, lower-grade tumor (TA2) [2] were analyzed. Thus, TA1 always represented the higher-grade nodule compared to TA2. Note, whereas each patient was assigned a patient-specific overall grade (*i*.*e*. ‘low’, ‘intermediate’ or ‘high’), each tumor area was additionally assigned an individual grade group based on its histological appearance. According to current ISUP guidelines, the grading of the entire prostate specimen depends on the size and grade of individual nodules [28]. Thus, it is possible that the patient grading is lower than the grading of the most aggressive nodule, if another lower-grade nodule is larger. Tumor regions contained at least 70% tumor cellularity and the distance between the analyzed areas (TA1 versus TA2) was at least 5 mm. Altogether, we obtained 105 prostate tissue specimens (**Additional file 2: Table S1**). Three adjacent tissue biopsies of the dimensions 0.6 × 0.6 × 3.0 mm were punched from each representative region for exome sequencing, CNA (derived from the exome sequencing data), RNA sequencing (RNA-seq), and quantitative proteomic analysis using the PCT-SWATH technology [23] respectively. Proteomic analysis was performed in duplicates for each tissue sample. Peripheral blood samples from each patient were also subjected to exome sequencing and served as the genomic wild-type reference (**Fig. 1**). All three types of grading (*i*.*e*. patient-specific overall grading, TA1 grading and TA2 grading) were predictive of the recurrence-free survival (RFS) in our study.

**Figure 1.**
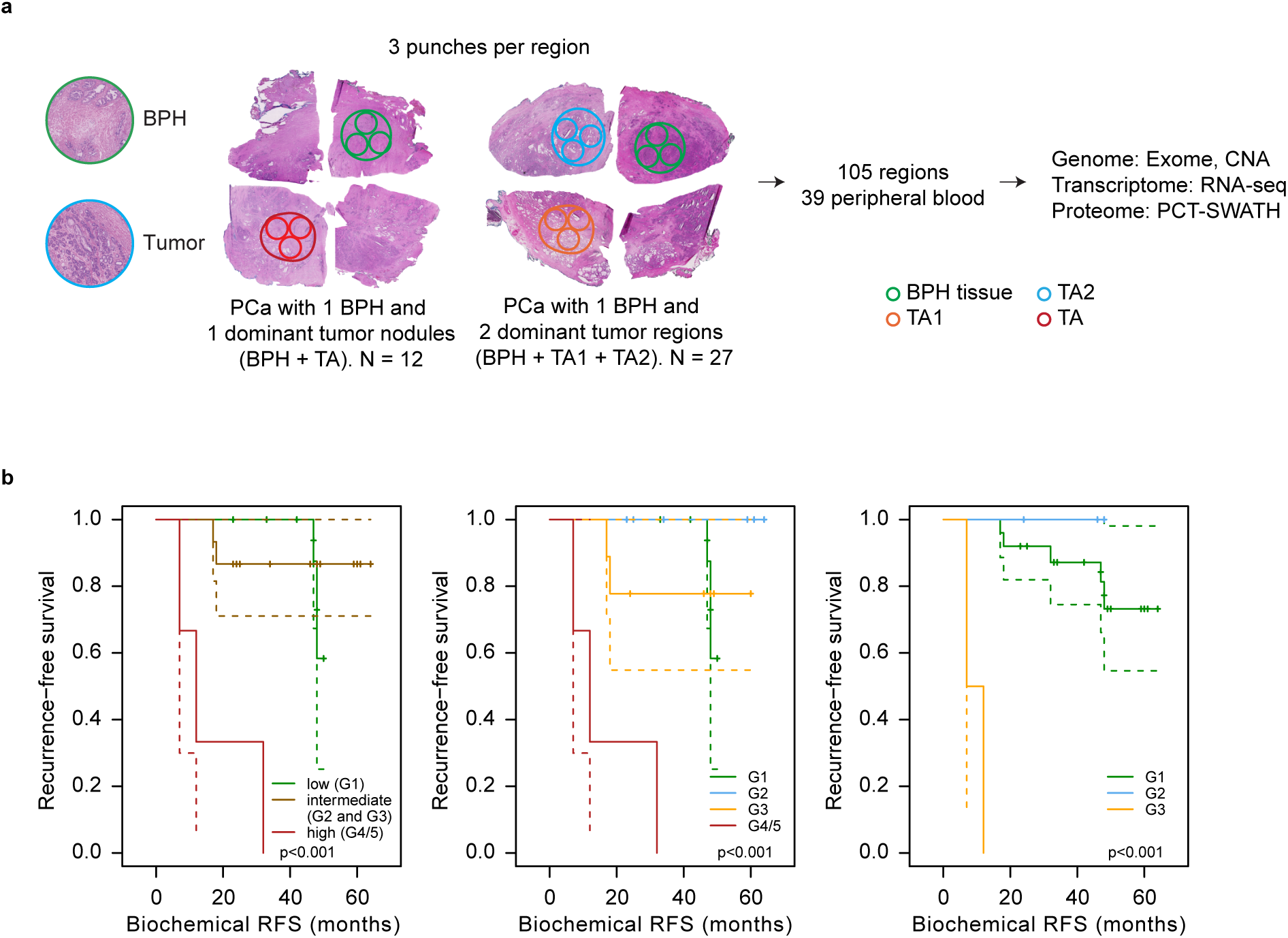
Proteogenomics analysis of 105 tissue regions from 39 PCa patients. **a** Representative immunohistochemistry images of prostate tissues and the selection of BPH and tumorous tissue regions for genome, transcriptome and proteome analysis. **b** Kaplan-Meier curves for our cohort when the patients are stratified by the overall grade (left), the TA1 or TA grade group (middle) and the TA2 or TA grade group (right). Point-wise 95% confidence bands are shown for the whole range of time values.

In agreement with prior reports, we observed relatively few recurrent point mutations across patients (**Additional file 1: Fig. S2, Additional file 3: Table S2**), but substantial CNAs (**Additional file 1: Figs. S3** and **S4, Additional file 4: Table S3**). Mutations in SPOP, FOXA1, and MED12 reported in independent cohorts [4-9] were confirmed in this cohort. In total, 1,110 genes showed copy number gains in at least five samples or copy number losses in at least five samples (see **Additional file 1: Supplementary Text** for details). **Additional file 1: Fig. S4** shows the CNA status of signature genes representing known areas of recurrent CNAs in PCa-split into fusion-partner and non-fusion-partner genes-for instance loss of PTEN and gain of MYC in high-grade PCa [29]. Likewise, our data confirmed the differential expression of several transcripts/proteins that had previously been suggested as PCa biomarkers or which are known oncogenes in other tumor types (**Additional file 1: Supplementary Text and Fig. S5, Additional file 5: Table S4** and **Additional file 6: Table S5**). We further identified somatic fusions from the RNA-seq data. A large fraction of the tumors harbored ETS family gene fusions, which are frequently detected in PCa [8, 9, 11]. ETS fusions were mutually exclusive and appeared in tumors from all grade groups (**Additional file 1: Fig. S6**; **Additional file 5: Table S4**). This consistency with previously published results confirmed the quality of our data and motivated us to go beyond previous work by performing a network-based multi-omics multi-gene analysis.

### Molecular perturbations correlate with tumor grade

Mutational burden is associated with PCa risk [8, 9, 11]. Hence, as a first step towards a cross-layer analysis, we asked if high-grade PCa would generally be affected by stronger alterations (compared to low-grade PCa) at the genome, transcriptome, and proteome layer [30]. For that purpose, we devised molecular perturbation scores that quantified the number of affected genes/proteins and the extent to which these genes/proteins were altered in the tumor specimens compared to their benign controls (see the ‘**Methods**’ section for details). In the case of the DNA layer these scores carry a similar meaning as established mutational burden scores. However, we wanted to capture effects at all three molecular layers measured in this study. Higher-grade tumors (G3 and G4/5) exhibited significantly higher molecular perturbation scores than lower-grade tumors (G1 and G2). Those differences were statistically significant in all but one case (*P* value < 0.05, one-sided Wilcoxon rank sum test, **Fig. 2**). The CNA perturbation magnitude exhibited the highest correlation with the PCa grading, confirming prior studies documenting the tight association between CNA, histopathological grade and risk of progression [4, 5, 31]. Further, we found that mRNA fold changes (FCs) correlated more strongly with CNAs of the same genes than protein FCs (average CNA-mRNA Spearman p = 0.1 and average CNA-protein Spearman p = 0.02). This observation is in agreement with previous work, which suggested that copy number changes are to some extent buffered at the protein level [17, 21, 32]. Interestingly, we observed that proteins known to be part of protein complexes were significantly less strongly correlated with the FCs of their coding mRNAs than proteins not known to be part of protein complexes (*P* value < 2.6e-11, one-sided t-test, **Additional file 1: Fig. S7**). This result is consistent with the concept that protein complex stoichiometry contributes to the buffering of mRNA changes at the level of proteins [21, 22, 33-35]. Thus, molecular patterns in high-grade PCa are more strongly perturbed at all layers and the effects of genomic variation are progressively but non-uniformly attenuated along the axis of gene expression.

**Figure 2.**
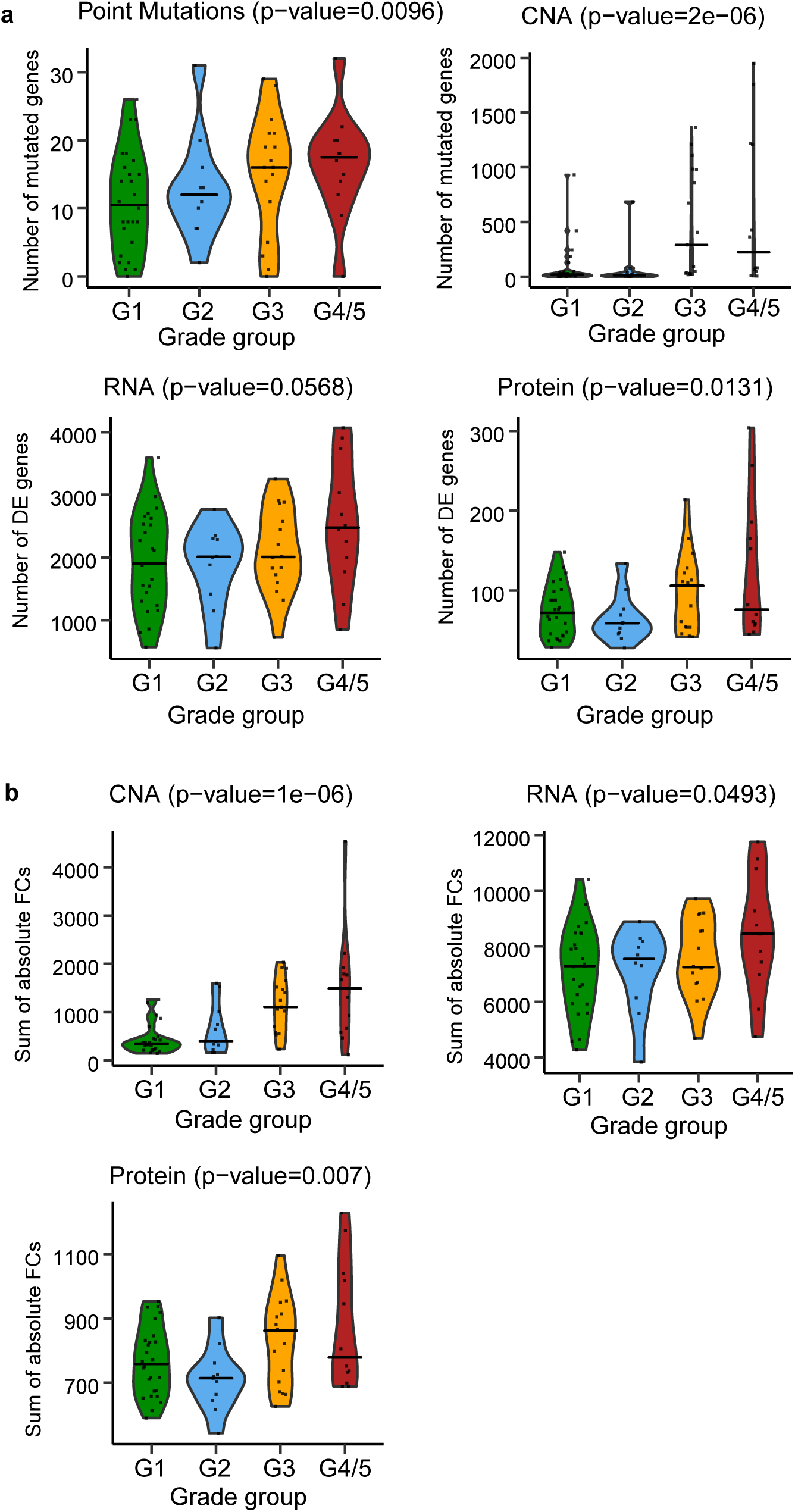
Molecular perturbation scores for point mutations, CNAs, transcriptome and proteome data. **a** Distributions of the first type of molecular perturbation scores (DE_count’s) for the four grade groups (visualized as violin plots) at the mutation layer (upper left), CNA layer (upper right), mRNA layer (lower left) and protein layer (lower right). Points represent the actual values. The horizontal lines correspond to the median value in each of the four grade groups. **b** Distributions of the second type of molecular perturbation scores (DE_sum’s) for the four grade groups (visualized as violin plots) at the CNA layer (upper left), mRNA layer (upper right) and protein layer (lower left). Points represent the actual values. The horizontal lines correspond to the median value in each of the four grade groups. *P* values (in each of the titles) show the significance of the one-sided Wilcoxon rank sum test where the values of G3 and G4/5 are gathered together and compared to the values of G1 and G2 (also gathered together).

### Effects of distinct CNAs converge on common proteins

It has previously been suggested that mutations affecting different genes could impact common molecular networks if the respective gene products interact at the molecular level [36]. However, previous analyses were mostly restricted to individual molecular layers. For example, it was shown that genes mutated in different patients often cluster together in molecular interaction networks [36]. Yet, effects of these mutations on transcript and protein levels remained unexplored in this case.

Previous work of Sinha and colleagues already suggested extensive *trans*-effects of CNAs on mRNA and protein levels [17]. Thus, here we aimed to systematically explore how different CNA events would impact on the level of one common protein. In order to prioritize potentially interesting proteins for such an analysis, we focused on the 20 proteins with the largest average absolute FCs across all tumor specimens (**Additional file 1: Fig. S8, Additional file 7: Table S6**). Thus, these proteins represent a set of proteins that was strongly affected across most tumors independent of tumor grade. Among them was PSA (KLK3), and several other well established PCa-associated proteins like AGR2 [37], MDH2 [38], MFAP4 [39] and FABP5 [40]. RABL3 was one of the most strongly down-regulated proteins, which is a surprising finding as RABL3 is known to be up-regulated in other solid tumors [41, 42]. Interestingly, in most cases these proteins were from loci that were not subject to CNAs (**Additional file 1: Fig. S8, Additional file 7: Table S6**), hinting that independent genomic events would impact on these target proteins *via* network effects in *trans*.

Among those top targets we selected AGR2, ACPP, POSTN and LGALS3BP for further analysis (**Fig. 3a**), because these proteins/genes had correlated protein-and mRNA FCs; thus, protein level changes were likely caused by cognate mRNA level changes. To identify potential regulators for each target gene, we used the STRING gene interaction network [43] and selected putative effectors at most one edge away from the target genes. Further, we required that neighbors or the target itself were subject to CNAs in at least four tumor samples (‘**Methods**’ section). By including the target itself we account for potential CNA *cis*-effects. However, only POSTN passed that filter. This filtering identified 13 neighbors of ACPP, 28 neighbors of POSTN (and POSTN itself), 14 neighbors of LGALS3BP and one neighbor for AGR2, which was not further considered. Next, we correlated CNAs of those neighbor genes with the mRNA FCs of the respective target genes (**Fig. 3b**). We then used the non-neighboring genes (*i*.*e*. the network complement) to generate a background distribution of CNA-target correlations specifically for each target. Here, we also only considered genes with at least four CNAs across the tumor samples. Since STRING reports predicted functional associations between genes we expected only a minority of the neighbors to actually correlate with their putative targets. Further note that edges in STRING could represent indirect gene-gene relationships. Yet, in the case of ACPP we found that CNA levels of its 13 neighbors were on average more strongly correlated with ACPP mRNA FCs than the complement (**Fig. 3b**). This observation does not preclude the possibility that also some of the POSTN and LGALS3BP neighbors impacted on their mRNA levels in *trans*. However, the fact that ACPP neighbors were on average more strongly correlated with ACPP mRNA FCs, suggested to us that multiple of its network neighbors might be involved in tumorigenic down-regulation of ACPP.

**Figure 3.**
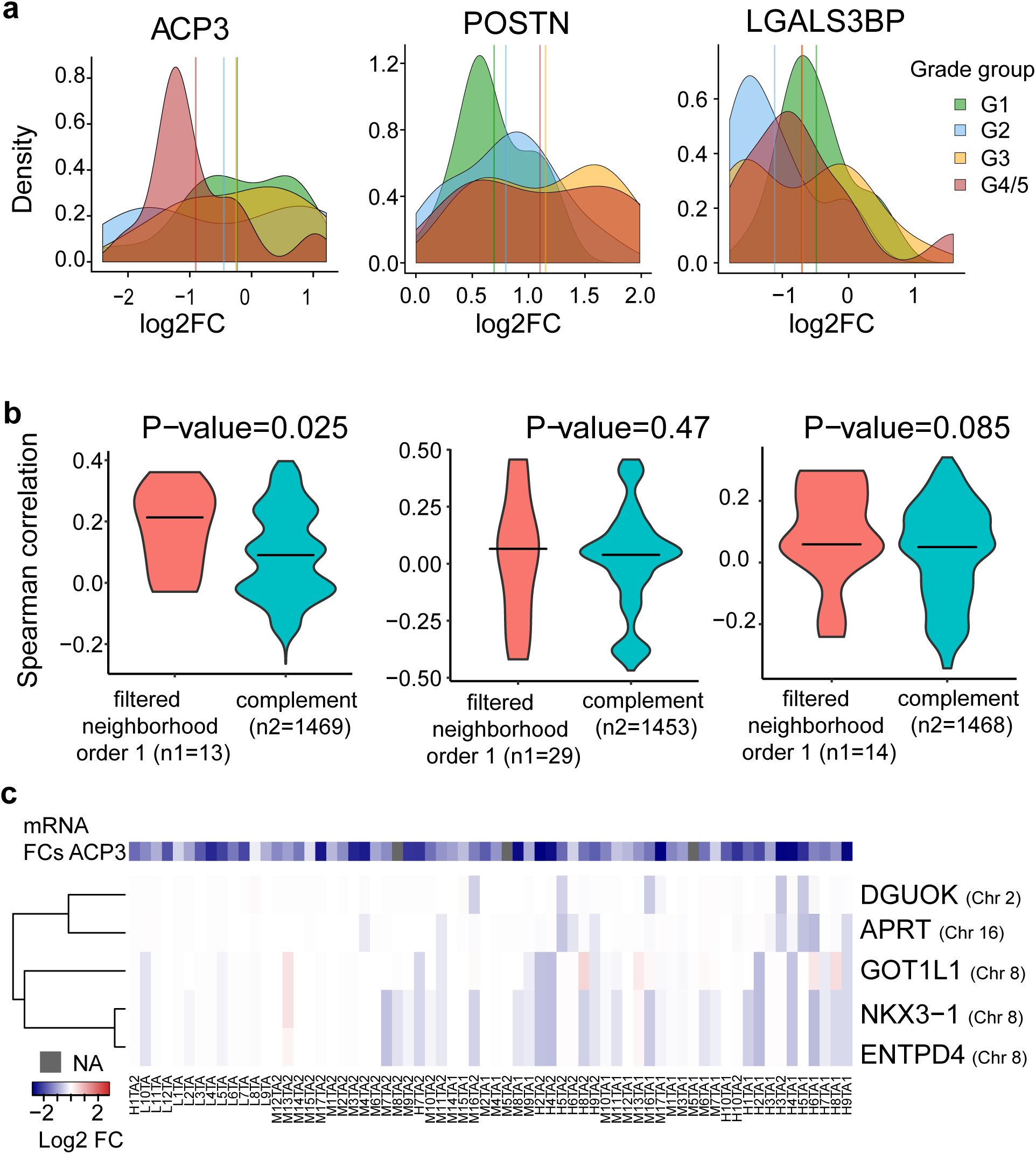
Target genes and putative effectors. **a** Density plots of the FCs in the four grade groups for three selected proteins (ACPP, POSTN, LGALS3BP) among the 20 highest scoring (score: mean of the absolute FCs across all tumor samples) proteins. Vertical lines correspond to the average FC in each of the four grade groups. These proteins were selected as target genes to identify potential regulators. **b** Distributions of the Spearman correlations of the mRNA target gene FCs with the CNAs of the ‘filtered neighborhood order one’ and the ‘complement’, for the three target genes. The first set/group consists of the target itself and of those neighbors in STRING with confidence above 0.2, while the second consists of the remaining network genes in STRING. Both sets are filtered out for genes subject to CNAs in less than four tumor samples. *P* values (in each of the titles) show the significance of the one-sided t-test. **c** Heatmap of the CNA matrix reduced to the significant regulators of the target gene ACPP output by the fitted elastic net model (*i*.*e*. those with a non-zero beta coefficient). The columns are ordered based on the grade group while there is a hierarchical clustering of the rows. The added colorbar depicts the mRNA FCs of the target gene ACPP.

ACPP, which is also known as ACP3 or PAcP, is a prostate-specific acid phosphatase with a critical role in PCa etiology and has been suggested as a PCa biomarker long before PSA [44]. ACP3 is known to inhibit cell proliferation and is therefore typically down-regulated in PCa [45], despite elevated ACP3 protein levels in patient blood [44]. In our cohort ACP3 levels were strongly down-regulated in all of the high-grade patients and in the vast majority of low-and intermediate-grade patients, suggesting that ACP3 down-regulation represents an early event during PCa evolution. Despite its established role in PCa, little is known about the oncogenic driver events down-regulating ACP3 [44].

We speculated that CNA events affecting different ACPP neighbors might be in operation in different tumor specimens. Thus, to further narrow the list of candidates we devised a multi-dimensional regression approach modeling the combined CNA effects of neighbors on ACPP mRNA FCs (**‘Methods’** section). Five neighbors (DGUOK, APRT, GOT1L1, NKX3-1 and ENTPD4) had statistically significant effects in that multi-dimensional model (**Fig. 3c**). We utilized two independent PCa cohorts (TCGA; [8] and MSKCC [31]) to validate potential effects of those five genes: we computed the association between the CNAs of each significant regulator and the corresponding mRNA log-FC of ACPP in each cohort and confirmed that the signs of the effects (i.e. effect directions) were the same for all five genes in all cohorts. Among those hits was NKX3-1, which is a prostate-specific tumor suppressor gene and loss of a single allele may predispose to prostate carcinogenesis [46, 47]. NKX3-1 is a transcription factor found to have substantial *tran*s-effects in PCa [17]. Consistent with its potential role as an ACPP regulator, NKX3-1 has been found to bind within 1 kb of the transcription start site of ACPP ([48]; GEO GSE40269). Interestingly, the CNA signatures of the five putative regulators split into two clusters affecting two distinct sets of patients (**Fig. 3c**): the first one harboring joint deletions of DGUOK and APRT, the second one harboring joint deletions of NKX3-1, ENTPD4 and GOT1L1. The latter three genes are all encoded on Chromosome 8 and thus, their deletion may be due to single CNA events. DGUOK and APRT however, are encoded on different chromosomes. Importantly, these events were clonal in most cases, *i*.*e*. they were mostly common to both tumor samples of a given patient. Hence, our network analysis hints that distinct deletions in the network vicinity of ACP3 can lead to the repression of this anti-proliferative protein. Taken together, these findings suggest that tumor mechanisms in different patients converged on common protein endpoints.

### Joint network effects of CNAs drive tumor progression

The analysis above identified molecular networks driving tumor alterations and thus indicated altered biochemical states that were common to most tumor specimens. To identify sub-networks that specifically distinguish high-grade from low-grade tumors, we performed a distinct network analysis: we mapped our data onto the STRING gene interaction network [43], and employed network propagation [49, 50] separately to the CNA, transcriptome and proteome data for each of the tumor samples. We excluded point mutations from this analysis as their frequency was too low in our cohort. By combining published molecular interactome data with a network propagation algorithm [36, 49], we aimed to ‘enrich’ network regions with many perturbed genes/proteins. We reasoned that the convergent consequences of genomic variants on common network regions would be indicative of specific biochemical functions that are important for the tumor biology. We therefore identified genes/proteins in network regions that showed a higher score (or a lower score) in high-grade (G4/5) relative to lower-grade (G1) tumor groups at all three levels (**Fig. 4a, b**; ‘**Methods**’ section). This analysis identified sub-networks consisting of over-and under-expressed genes (relative to the benign controls). We found 57 amplified genes (**Additional file 7: Table S6**) for which transcripts and proteins were often over-expressed in high-grade PCa (**Fig. 4a**) and 21 genes with copy number loss (**Additional file 7: Table S6**) for which transcripts and proteins were often down-regulated compared to lower-grade tumors (**Fig. 4b**).

**Figure 4.**
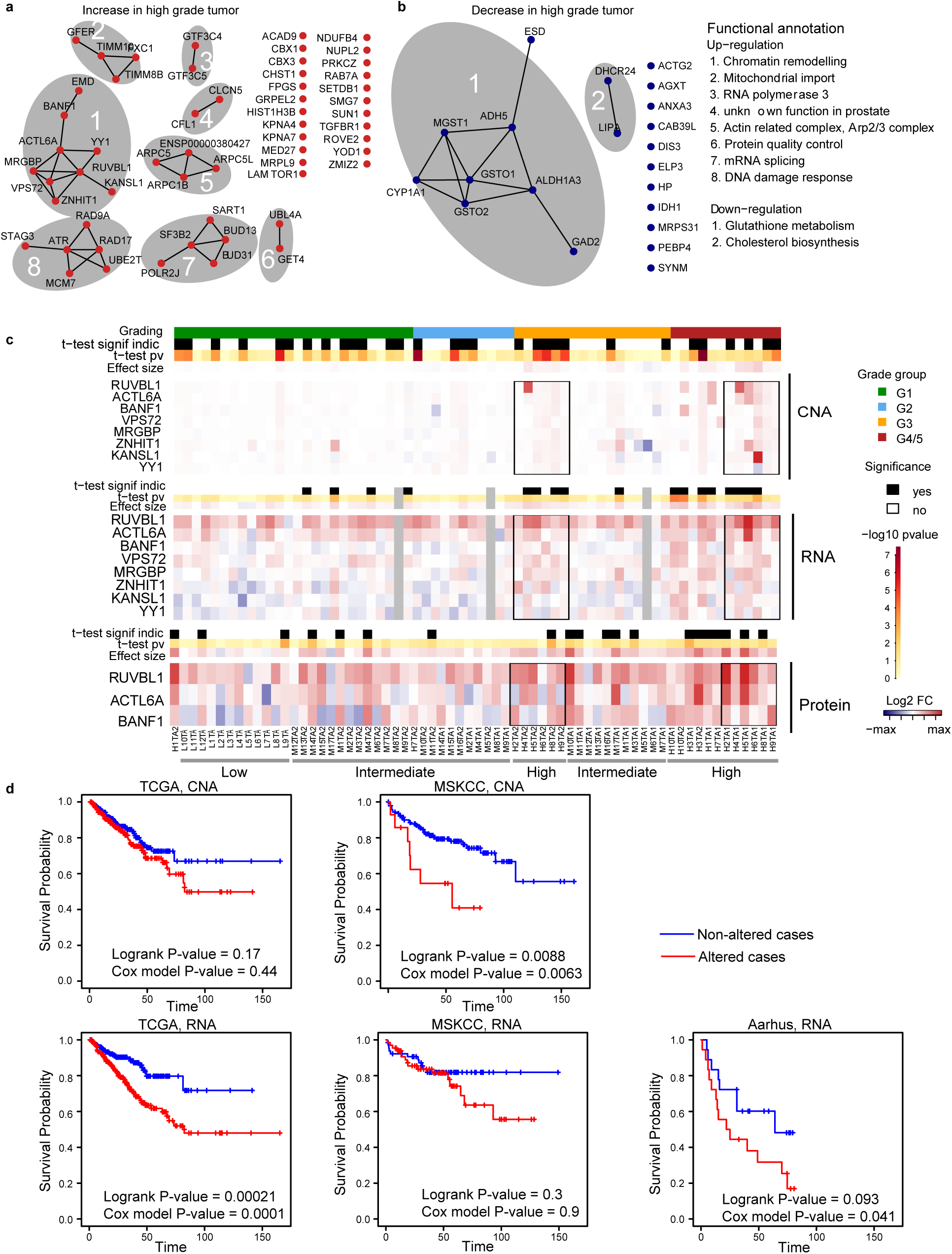
Cross-omics networks distinguishing high-grade from low-grade tumors. **a** Sub-networks consistently up-regulated in high-grade (G4/5) compared to low-grade (G1) tumors across all three layers (CNA, mRNA and protein). **b** Same as in (a) but down-regulated genes. Functional annotation of the sub-networks in (a) and (b) with more than one node is given. All edges in (a) and (b) are supported by either experimental or database evidence (STRING evidence ≥ 0.348). **c** CNA, RNA and protein FCs of Network Component 1 from (a). Samples are ordered by grade group (top bar). T-test results comparing Network Component 1 members against no change (*i*.*e*. 0) are shown for each molecular layer along with the average FC across Network Component 1 members (‘effect size’). Black box marks selected matching samples from patients with G4/5 and G3 tumor areas; i.e. tumor sample pairs from identical patients. Those areas exhibit weak, but common amplifications of Network Component 1 members at the CNA and RNA layers. mRNA samples in grey were removed due to low RNA quality. Gray bars at the bottom show the grade group of the patients (low, intermediate, high) where the samples have (mainly) come from. **d** Kaplan-Meier curves for ‘altered’ and ‘unaltered’ samples, where ‘altered’ is defined as an effect size greater or equal to the median effect. Results for three independent studies, TCGA (left), MSKCC (middle) and Aarhus (right) using the corresponding CNA data when available (first row) and mRNA data (second row). The Cox model *P* value corresponds to the *P* value of the variable of interest (*i*.*e*. average copy number change (CNA) or average z-score (mRNA) of Network Component 1) from the fitted Cox model after adjusting for patient age (when available, *i*.*e*. for TCGA and Aarhus).

Among the up-regulated network nodes, we observed genes modulating the stability of chromatin, such as chromatin-binding protein Chromobox 1 (CBX1) [51], SET Domain Bifurcated 1 (SETDB1) [52], a function linking to H3K27me3 and H3K9me3 in chromatin, and CBX3 (known as HP1-γ) [53]. SETDB1 is an oncogene in melanoma [54] and has also been found to be over-expressed in PCa and cell lines [55]. Further, we found genes involved in DNA damage repair, such as SMG7 [56] and ATR [57], and PRKCZ [58], which had already been suggested as a biomarker prognostic for survival in PCa [59]. Multiple actin related proteins including ARPC1B [60], ARPC5 [61], ACTL6A [62], and CFL1 [63], which are markers for aggressive cancers, were part of the up-regulated network nodes. Moreover, the up-regulated genes contained proteins related to the cell cycle like BANF1 and proteins interacting with the centrosome including LAMTOR1 and RAB7A that had already been associated with PCa [64]. Finally, several signaling molecules with known roles in PCa were up-regulated, such as the transcription factor Yin Yang 1 (YY1) [65], the TGF-β receptor TGFBR1 [66], and KPNA4, which promotes metastasis through activation of NF-κB and Notch signaling [67]. Thus, up-regulated network nodes are involved in DNA/chromatin integrity and growth control.

Likewise, several of the down-regulated genes had functions associated with PCa. For example, the oxidative stress related gene MGST1, which is recurrently deleted in PCa [68]. ALDH1A3 is a direct androgen-responsive gene, which encodes NAD-dependent aldehyde dehydrogenase [69]. DHCR24 is involved in cholesterol biosynthesis and regulated by the androgen receptor [70]. Polymorphisms in CYP1A1 are associated with PCa risk in several meta-analyses among different ethnicities [71-73].

Further, our network analysis is suggesting tumor mechanisms converging on genes that are known contributors to PCa tumor biology. For example, the PCa-associated gene SF3B2 [74, 75] was only weakly amplified in some of the high-grade tumors (average log_2_FC = 0.016) and mRNA levels showed similarly small changes (average log_2_FC = 0.024). On the other hand, the SF3B2 protein levels were consistently and more strongly up-regulated across tumors (average log_2_FC = 0.31), especially within the high-grade tumors (**Additional file 1: Fig. S9**). Another example is UBE2T whose over-expression is known to be associated with PCa [76]. Unfortunately, we could not quantify the corresponding protein levels. However, we observed a strong and consistent mRNA over-expression across several tumors (average log_2_FC = 0.73), even though at the DNA level the gene was only weakly amplified (average log_2_FC = 0.023; **Additional file 1: Fig. S9**). Our findings of more heterogeneous CNAs, but more uniform mRNA and protein alterations point on convergent evolutionary mechanisms, as we move along the axis of gene expression.

Next, we analyzed the largest connected component with genes up-regulated in advanced disease in more detail (see the ‘**Methods**’ section). It consists of the nine nodes EMD, BANF1, ACTL6A, YY1, RUVBL1, KANSL1, MRGBP, VPS72 and ZNHIT1 (**Fig. 4a**), and is referred to in the following as Network Component 1 (**Additional file 7: Table S6**). Seven of these proteins are involved in chromosome organization which may induce genomic alterations and influence the outcome of multiple cancers including PCa [77]. For example, the actin-related protein ACTL6A is a member of the SWI/SNF (BAF) chromatin remodeling complex[78], and a known oncogene and a prognostic biomarker for PCa [79]. Further, ACTL6A, RUVBL1 and MRGBP are together part of the NuA4/Tip60-HAT complex, which is another chromatin remodeling complex involved in DNA repair [80]. Likewise, KANSL1 is involved in histone post-translation modifications, while VPS72 is a member of histone-and chromatin remodeling complexes [81]. Thus, Network Component 1 consists of genes involved in chromatin remodeling and DNA repair, many of which are known to be involved in cancers.

Several samples were characterized by a small, but consistent DNA amplification of multiple members of Network Component 1 (**Fig. 4c**). Out of the 66 tumor samples, there were 30 samples – belonging to all grade groups – with a weak but remarkably consistent DNA amplification of Network Component 1 members, while the high-grade samples had stronger amplifications on average (*i*.*e*. larger effect sizes). Importantly, gene members of Network Component 1 were dispersed across eight chromosomes (**Additional file 7: Table S6**). The parallel DNA amplification of these genes is therefore the result of multiple independent CNA events, while the signal on any single gene alone was too weak to be significant in isolation. Further, members of Network Component 1 were consistently amplified in both tumor areas (i.e. TA1 and TA2) of six patients (H2, H4, H5, H6, H8, and H9; **Fig. 4c**), thus establishing them as likely clonal events. In some but not all cases, the amplifications led to a small, but consistent increase in mRNA expression of the amplified gene loci (**Fig. 4c**). We were able to reconcile 40 tumor samples with a significant enrichment of this network component in either the CNA or mRNA layer. Unfortunately, only three out of the nine proteins were detected in our proteomics experiments (**Fig. 4c**). Interestingly, patients where the DNA amplifications led to transcript over-expression were almost always high-grade patients, whereas patients where the amplification affected gene expression to a smaller extent were low-or intermediate-grade patients (**Fig. 4c**). Further, we noticed that TA2 samples graded as G3 from high-grade patients carried amplifications of Network Component 1, whereas tumor areas graded as G3 from intermediate-grade patients did not have amplifications of this network component (**Fig. 4c**). Thus, although the tumor areas were histologically equally classified, tumor areas from high-grade patients carried a CNA signature and expression patterns reminiscent of the high-grade areas from the same patients. Therefore, within the cohort tested the joint DNA amplification of this network component along with RNA up-regulation is a signature of high-grade tumors. Curiously, the higher-grade tumor areas of those high-grades patients (TA1) carried stronger DNA amplifications than the respective lower-grade areas (TA2), which implies that the progressive amplification of Network Component 1 during tumor evolution may contribute to an increasingly aggressive phenotype. To further corroborate the clinical relevance of this network perturbation, we analyzed published datasets of three additional PCa cohorts (TCGA[8], MSKCC [31], and Aarhus [82]), together comprising a total of 709 patients with known clinical outcome. We found that amplification of genes from Network Component 1 was a significant predictor of reduced RFS in the MSKCC cohort (*P* value = 8.8e-3, log-rank test). In the TCGA cohort, we observed the same trend although the difference in RFS was not statistically significant (*P* value = 0.17; **Fig. 4d**). Additionally, we found that over-expression of genes from Network Component 1 was a significant predictor of reduced RFS in the TCGA cohort (*P* value = 2.1e-4, log-rank test), which was the cohort with the largest number of patients. In the other two cohorts we observed the same trend, although the difference in RFS was not statistically significant (*P* value = 0.30 and 0.093 for MSKCC, and Aarhus; **Fig. 4d**). Thus, both CNA and RNA changes of Network Component 1 are predictive of the time to relapse in independent cohorts. To also account for covariates, we fitted Cox proportional-hazards models with the age and the copy number burden as (additional) covariates. When including only the age in our model, the results showed minor changes (**Fig. 4d**). When including both the age and the copy number burden in our model, the effect direction of Network Component 1 remained the same but was not statistically significant anymore (*P* value = 0.055 for TCGA, mRNA and *P* value = 0.064 for MSKCC, CNA).

In conclusion, our findings suggest that relatively weak but broad CNAs of entire network components are associated with high-grade tumors and that the presence of some of these perturbations in lower-grade tumors may be predictive of the future development of a more aggressive phenotype.

### Analysis of distinct tumor nodules defines intra-patient heterogeneity (TA1 versus TA2 comparison)

The CNA patterns (**Additional file 1: Fig. S4**) and the Network Component 1 analysis (**Fig. 4c**) suggest that different tumor areas from the same patient shared several mutations. Such common signatures are expected if different tumor nodules originate from a common clone. If this was true, we would expect mutational signatures to be more similar between different nodules from the same patient than between patients, even though mutated genes may be shared across patients. To compare the intra-and inter-patient molecular heterogeneity at the levels of CNAs, transcript, and protein FCs, we computed the Pearson correlation between tumor area 1 (TA1) and its paired tumor area 2 (TA2) for each layer and all of the 27 patients with two characterized tumor areas (25 for the mRNA, see the ‘**Methods**’ section and **Additional file 1: Supplementary Text**). As a control, we also computed all pairwise Pearson correlations between the samples within each of the grade groups (*i*.*e*. inter-patient correlation). As expected, paired TA1 and TA2 from the same patient were on average more strongly correlated to each other compared to samples from different patients within the same grade group. This finding was consistent for all omics layers (**Fig. 5a**), and was more pronounced at the CNA and mRNA layers compared to the protein layer.

**Figure 5.**
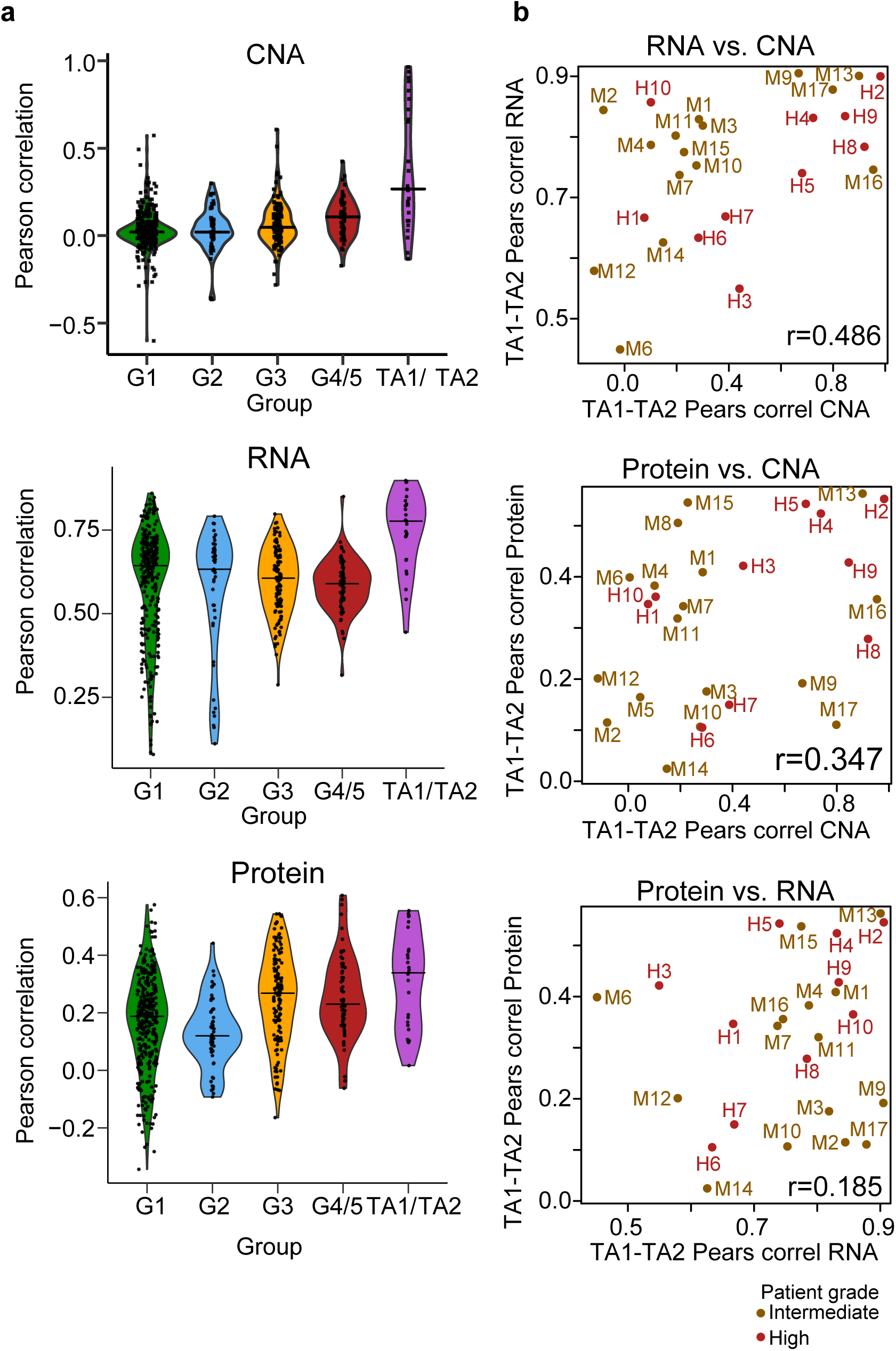
Within-patient similarity at the different layers. **a** Distributions of the within-group similarities for the four grade groups (visualized as violin plots) based on the Pearson correlation at the CNA layer (upper), mRNA layer (middle) and protein layer (lower). A ‘violin’ with the correlations between TA1 and paired TA2 for the different patients has been added to all three plots and colored in purple. Points represent the actual values. The horizontal lines correspond to the median value in each of the groups. *P* values from the one-sided Wilcoxon rank sum test comparing the within-patient to the within-group similarities (where all values from the four groups are gathered together): 8.97e-09 for the CNA, 4.42e-08 for the mRNA and 6.27e-04 for the protein layer. **b** The correlations between TA1 and paired TA2 for the different patients at one layer are plotted against the corresponding correlations at another layer for each pair of layers: mRNA versus CNA (upper), protein versus CNA (middle) and protein versus mRNA (lower). The points are labeled and colored based on the overall grade in all plots; r: Pearson correlation.

Next, we tested whether a high correlation at the level of CNAs also implies a high correlation at the level of mRNA and proteins. We tested this idea by ‘correlating the correlations’, *i*.*e*. we correlated the TA1-TA2 correlation of CNA profiles with the correlation between the mRNA and protein profiles of the same tumor areas (**Fig. 5b**). Indeed, a higher correlation of two tumor areas at the level of CNA correlated significantly with a higher correlation at the level of mRNA (r=0.49, *P* value=0.014). In other words, knowing how similar two tumor areas of a patient are at the CNA level supports a prediction of their similarity at the mRNA level (and conversely). Although the correlation between protein and CNA was not statistically significant, it followed the same trend (r=0.35, *P* value=0.076).

Comparing molecular similarity across omics layers allowed us to identify specific types of patients. The patients H2, H4, M13 had highly correlated tumor areas at all three layers (upper right corner in all scatterplots of **Fig. 5b**). Likely, the tumor areas of these patients have a common clonal origin (**Additional file 1: Fig. S3**). In contrast, patients M12 and M14 had weakly correlated tumor areas at all levels (bottom left corner in all scatterplots of **Fig. 5b**). These tumor nodules either have independent clonal origins or they diverged at an earlier stage during tumor evolution (**Additional file 1: Fig. S3**) [16]. For example, in the case of patient M12 large parts of the genome were not affected by CNAs in the benign sample as well as in TA1 and TA2. However, as shown on **Additional file 1: Fig. S3**, a large region was amplified in TA1, whereas the same region was deleted in TA2. This is consistent with a scenario in which TA1 and TA2 show parallel evolution. A third class of patients is exemplified by the patients M9 and M17, who showed a high correlation between their tumor areas on the CNA and mRNA levels, but not on the protein level. Yet other patterns were apparent in patients M4, M7, and H10. They showed similar mRNA and protein patterns in the two tumor areas, but relatively uncorrelated CNAs. The results here apply to global proteome patterns and therefore hint that such convergent network effects of CNAs can be frequent. We confirmed that protein-level similarity correlated with similar histological characteristics of the tumor areas. **Additional file 1: Fig. S10** shows formalin-fixed paraffin-embedded (FFPE) tissue microarray images (duplicates) from the analyzed tumor nodules (TA1 and TA2, diameter 0.6 mm), further underlining the hypothesis that ultimately protein-level alterations are responsible for common cellular phenotypes. Although we cannot fully exclude the possibility that some of these results were affected by technical noise in the data, our findings suggest that transcript alterations can frequently be buffered at the level of proteins (patients M9, M17, **Additional file 1: Fig. S7**) and that convergent evolutionary processes may lead to the alteration of common proteins (patients M4, M7, H10). We also note that our findings are specific to the two tumor areas available in this study and could be different if other nodules had been sampled for each of the patients. However, our findings on patients with weakly correlated tumor areas at all levels like M12 and M14 suggest that these patients might carry more than one disease [16].

## Discussion

Despite twenty years of oncological research involving genome-scale (omics) technologies, we know remarkably little about how the discovered genomic alterations affect the biochemical state of a cell and consequently the disease phenotype. In particular, little is known about how genomic alterations propagate along the axis of gene expression [17, 18]. Here, we have exploited recent technological advances in data acquisition that made it possible to characterize small samples of the same tumor specimens at the level of genomes, transcriptomes, and proteomes and advances in computational strategies towards the network-based integration of multi-omics data.

In our study, samples were generated from small, less than 1 mm diameter punches in immediate spatial proximity in the tumor and subsequently profiled at all three ‘omics layers’ (DNA, RNA, proteome). Due to the large spatial heterogeneity of PCa [14, 24], this design - which is so far uncommon for studies profiling multiple layers from tumor specimens - was instrumental for increasing the comparability of the various omics layers and thus facilitated the analysis of molecular mechanisms. Our key findings are: (1) we confirmed the importance of CNAs for PCa biology and the alteration of many known PCa-associated genes at the transcript-and protein-level; (2) we revealed a generally elevated molecular alteration of high-grade tumors compared to lower-grade tumors; (3) although our study confirmed large within-and between-patient genomic heterogeneity, (4) we detected molecular networks that were commonly altered at the mRNA and protein-level. The fact that many of those target molecules are known drivers of PCa tumorigenesis, supports the notion that these proteins/transcripts are subject to convergent evolutionary mechanisms.

We integrated the three omics layers using a network-based approach as opposed to directly comparing gene perturbations (mutations) to gene products (transcripts and proteins). Using genome data only, it had previously been hypothesized that whereas the identity of specific mutated genes may differ between tumors, those mutations might still affect common molecular networks [36]. In other words, tumor phenotypes are determined by the perturbation of molecular networks and not by the perturbation of isolated genes. Our study provides experimental evidence that such network effects are indeed propagated to subsequent molecular layers and that this effect propagation may be clinically relevant. A very prominent example is the indication derived from our data that the long-known PCa gene ACPP (ACP3) is downregulated through diverse CNA events. Of particular interest is the potential role of NKX3-1 in ACP3 downregulation. Although both genes have a well-established PCa association their regulatory relationship had not been reported so far (to our knowledge).

Our multi-omics network analysis revealed molecular sub-networks that distinguished high-grade PCa tumors from low-grade tumors. Specifically, our analysis led to the identification of Network Component 1, a sub-network involved in chromatin remodeling and consisting of genes that were weakly amplified in intermediate-grade (G3) tumor specimens. Signals of individual gene members of this component were virtually indistinguishable from noise in our cohort. However, their consistent alterations across the network region, across molecular layers and the fact that the same genes showed enhanced signals in high-grade specimens, rendered this component highly interesting. The fact that copy number and expression changes of Network Component 1 members were predictive for survival in independent cohorts further supports the potential clinical relevance of this sub-network. Amplification of Network Component 1 was to some extent confounded with overall CNA burden (r=0.58 (TCGA, CNA), r=0.55 (TCGA, mRNA), r=0.34 (MSKCC, CNA), r=0.16 (MSKCC, mRNA)). However, the amplifications of Network Component 1 members were highly correlated and on average above the background of CNAs. Thus, the coordinated amplification of Network Component 1 does not simply mirror overall CNA burden. Our network-based cross-omics analysis identified nine other network components (**Fig. 4**) successfully capturing several known and potentially new PCa-associated genes. However, neither Network Component 1 nor any of the other network components was uniformly subject to CNAs across all high-grade patients. Instead, we found different network components modified in different patients and these sub-networks were involved in cellular processes as diverse as actin remodeling, DNA damage response, and metabolic functions, all of which are known contributors to PCa biology. This further underlines the large inter-patient variability of PCa and it demonstrates the diversity of molecular mechanisms leading to histologically similar phenotypes. Future prediction models of PCa including the ISUP grade groups, PSA levels and clinical stage might be improved by exploiting multi-omics network analyses. Detecting aggressive networks alterations in prostate biopsies would help clinicians to advice either active surveillance or active therapy. However, the development of such multi-dimensional biomarkers would require much larger patient cohorts.

Another distinguishing feature of this study was the simultaneous profiling of two different tumor regions in 27 out of the 39 patients. The profiling of multiple tumor regions from the same prostate helped to further highlight the enormous heterogeneity of PCa within patients and provided important insights into PCa evolution. The fact that Network Component 1 was more strongly affected in the paired higher-grade nodules of high-grade patients suggests that at least certain sub-networks are subject to an evolutionary process, that progressively ‘moves’ protein levels towards a more aggressive state. Generally, and at all molecular layers tested, the two paired tumor areas were more similar to each other compared to two samples from the same grade group but different patients, suggesting common evolutionary origins. Although the two tumor areas seemed to mostly originate from the same clone, this was not always the case. In some patients, different nodules exhibited different molecular patterns at all omics layers, suggesting early evolutionary separation. Thus, for the first time, current diagnostic, expert-level consensus guidelines [28] are supported by detailed proteogenomic data. Our findings support earlier claims that clonality itself might be a prognostic marker with implications for future, more tumor-specific treatment when targeted therapies become available also for PCa [16, 83].

Our study shows that all three molecular layers (genome, transcriptome and proteome) contributed valuable information for understanding the biology of PCa. In particular the DNA layer informed about causal events, clonality, and genomic similarity between tumors. The transcriptome was relevant for understanding the transmission of CNA effects to proteins and served as a surrogate in cases where protein levels remained undetected. The proteome was crucial for revealing protein-level buffering of CNA effects as well as for indicating convergent evolution on functional endpoints. In a routine diagnostic context though, measuring all three layers may not be feasible for the near future due to resource and time limitations. Thus, the identification of improved, routine-usable molecular markers for PCa diagnostics and prognosis remains an open problem [17].

## Conclusions

This study uncovered molecular networks with remarkably convergent alterations across tumor sites and patients. In particular, we identified a sub-network consisting of nine genes whose joint activity positively correlated with increasingly aggressive tumor phenotypes. The fact that this sub-network was predictive for survival in independent cohorts further supports its potential clinical relevance. At the same time though, our study also exposed a diversity of network effects: we could not identify a single sub-network that was perturbed in all high-grade tumor regions, let alone the observed distinct intra-patient alterations at all omics layers for some patients. Overall, our study has significantly expanded our understanding of PCa biology and serves as a model for future work aiming to explore network effects of mutations with an integrated multi-omics approach.

## Methods

### Patients and samples

A total of 39 men with localized PCa who were scheduled for RP were selected from a cohort of 1,200 patients within the ProCOC study and processed at the Department of Pathology and Molecular Pathology, University Hospital Zurich, Switzerland [25]. Each of the selected intermediate-and high-grade patients had two different tumor nodules with different ISUP grade groups. H&E (Hematoxylin and Eosin)-stained fresh frozen tissue sections of 105 selected BPH and tumor regions were evaluated by two experienced pathologists (PJW, NJR) to assign malignancy, tumor stage, and Grade Group according to the International Union Against Cancer (UICC) and WHO/ISUP criteria. This study was approved by the Cantonal Ethics Committee of Zurich (KEK-ZH-No. 2008-0040), the associated methods were carried out in accordance with the approved guidelines, and each patient has signed an informed consent form. Patients were followed up on a regular basis (every three months in the first year and at least annually thereafter) or on an individual basis depending on the disease course in the following years. The RFS was calculated with a biochemical recurrence (BCR) defined as a PSA ≥0.1 ng/ml. Patients were censored if lost to follow-up or event-free at their most recent clinic visit. Patients with a postoperative PSA persistence or without distinct follow-up data for the endpoint BCR were excluded from the analysis of BCR.

### Exome sequencing and somatic variant analysis

The exome sequencing (exome-seq) was performed using the Agilent Sure Select Exome platform for library construction and Illumina HiSeq 2500 for sequencing read generation. We mapped and processed the reads using a pipeline based on bowtie2 [84] (1.1.1) and the Genome Analysis Tools Kit (GATK) [85] (3.2-2). We detected and reported nonsynonymous variants or variants causing splicing changes using Strelka (1.0.14) and Mutect (1.1.7) combined with post-processing by the CLC Genomics Workbench (8.0.3). In this process, all tissue samples of a patient were compared to the respective blood sample.

Trimmomatic [86] (0.36) was used for adaptor clipping and low-quality subsequence trimming of the FASTQ files. Subsequently, single reads were aligned to the hg19 reference genome with bowtie2 with options “--very-sensitive -k 20”. We applied samtools [87] (0.1.19) and picard-tools (1.119) to sort the resulting bam files in coordinate order, merge different lanes, filter out all non-primary alignments, and remove PCR duplicates. Quality of the runs was checked using a combination of BEDtools [88] (2.21), samtools, R (3.1) and FastQC (0.11.2).

Bam files containing the mapped reads were preprocessed in the following way: indel information was used to realign individual reads using the RealignerTargetCreator and IndelRealigner option of the GATK. Mate-pair information between mates was verified and fixed using Picard tools and single bases were recalibrated using GATK’s BaseRecalibrator. After preprocessing, variant calling was carried out by comparing benign or tumor prostate tissue samples with matched blood samples using the programs MuTect [89] and Strelka [90] independently. Somatic variants that were only detected by one of the two programs were filtered out using CLC Genomics Workbench. So were those that had an entry in the dbSNP [91] common database and those that represented synonymous variants without predicted effects on splicing.

### CNA analysis of exome-seq data

The Bam files generated during the process of somatic variant calling were processed with the CopywriteR package (v.2.2.0) for the R software [92]. CopywriteR makes use of so-called “off-target” reads, *i*.*e*. reads that cover areas outside of the exon amplicons. “Off-target” reads are produced due to inefficient enrichment strategies. In our case on average 28.5% of the total reads were not on target. Briefly, CopywriteR removes low quality and anomalous read pairs, then peaks are called in the respective blood reference, and all reads in this region are discarded. After mapping the reads into bins, those peak regions, in which reads had been removed, were compensated for. Additionally, read counts are corrected based on mappability and GC-content. Finally, a circular binary segmentation is carried out and for each segment the log count ratios between tissue samples and the respective blood sample are reported as copy number gain or loss. The copy number of each gene in each sample was reported based on the log count ratio of the respective segment in which the gene was located. The overall performance of this CNA-calling approach was evaluated by comparing the results of the TA1 (and TA) samples with CNA results obtained by applying the OncoScan Microarray pipeline to FFPE samples from the same tumors (**Additional file 1: Fig. S11**).

### OncoScan Microarrays

OncoScan copy number assays were carried out and analyzed as described previously [93]. Briefly, DNA was extracted from punches of FFPE cancer tissue blocks. Locus-specific molecular inversion probes were hybridized to complementary DNA and gaps were filled in a nucleotide-specific manner.

After amplification and cleavage of the probes, the probes were hybridized to the OncoScan assay arrays. Scanning the fluorescence intensity and subsequent data processing using the Affymetrix® GeneChip® Command Console and BioDiscovery Nexus express resulted in log intensity ratio data (sample versus Affymetrix reference) and virtual segmentation of the genome into areas with copy number gain, loss or stability.

### RNA Sequencing

RNA sequencing was performed at the Functional Genomics Center Zurich. RNA-seq libraries were generated using the TruSeq RNA stranded kit with PolyA enrichment (Illumina, San Diego, CA, USA). Libraries were sequenced with 2×126bp paired-end on an Illumina HiSeq 2500 with an average of 105.2 mio reads per sample.

Paired-end reads were mapped to the human reference genome (GRCh37) using the STAR aligner (version 2.4.2a) [94]. Quality control of the resulting bam files using QoRTs [95] and mRIN [96] showed strong RNA degradation[97] in a significant fraction of the samples: mRIN classified 31 samples as highly degraded (**Additional file 1: Fig. S12, Additional file 5: Table S4**). In order to correct for this 3’ bias, 3 tag counting was performed as described by Sigurgeirsson et al [98] using a tag length of 1,000. After 3’ bias correction, three samples still showed a clear 3’ bias: the two tumor regions (TA1 and TA2) of the patient M5 and TA2 from patient M8 (**Additional file 1: Fig. S12**). These samples were excluded from subsequent analyses. Additionally, the BPH region of the patient M5 was excluded due to the exclusion of both its tumor regions.

FeatureCounts [99] was used to determine read counts for all genes annotated in ENSEMBL v75. Genes for which no read was observed in any of the samples in the original data were excluded from the analysis. Further, after 3 tag counting, all genes with without at least 1 read per million in N of the samples were removed. We chose N to be 10 which corresponds to the size of the smallest grade group (G2). In a last reduction step, all genes with more than one transcript were excluded, yielding a final set of 14,281 genes.

Read count normalization and differential gene expression analysis was performed using the R packages sva [100] and DESeq2 [101]. All benign tissues were considered biological replicates and differential gene expression for the individual tumor samples was determined against all benign tissues. Gene expression changes with an adjusted *P* value < 0.1 were considered significant.

### RNA-seq - 3’ bias correction

The 3 tag counting approach for 3’ bias correction was used on the RNA-seq dataset [98]. This approach requires changing of the annotation file in two steps: 1) isoform filtering and 2) transcript length restriction. As proposed in [98] for each gene we determined the highest expressed isoform within a set of high quality samples. As high quality samples we used all samples with an mRIN score greater than or equal to 0.02. This set contains 7 benign and 15 tumor samples. Isoform expression was determined using cufflinks [102]. As transcript length we chose 1,000bp.

### Gene fusions

FusionCatcher (version 0.99.5a beta) was used to determine gene fusions for all samples. Fusions classified as “probably false positive” are discarded unless they are also classified as “known fusion”.

### PCT assisted sample preparation for SWATH-MS

We first washed each tissue sample to remove O.C.T., followed by PCT-assisted tissue lysis and protein digestion, and SWATH-MS analysis, as described previously [23]. Briefly, a series of ethanol solutions were used to wash the tissues each tissue, including 70% ethanol / 30% water (30 s), water (30 s), 70% ethanol / 30% water (5 min, twice), 85% ethanol / 15% water (5 min, twice), and 100% ethanol (5 min, twice). Subsequently, the tissue punches were lysed in PCT-MicroTubes with PCT-MicroPestle [103] with 30 µl lysis buffer containing 8 M urea, 0.1 M ammonium bicarbonate, Complete protease inhibitor cocktail (Roche) and PhosSTOP phosphatase inhibitor cocktail (Roche) using a barocycler (model NEP2320-45k, PressureBioSciences, South Easton, MA). The lysis was performed with 60 cycles of high pressure (45,000 p.s.i., 50 s per cycle) and ambient pressure (14.7 p.s.i., 10 s per cycle). The extracted proteins were then reduced and alkylated prior to lys-C and trypsin-mediated proteolysis under pressure cycling. Lys-C (Wako; enzyme-to-substrate ratio, 1:40) -mediated proteolysis was performed using 45 cycles of pressure alternation (20,000 p.s.i. for 50 s per cycle and 14.7 p.s.i. for 10 s per cycle), followed by trypsin (Promega; enzyme-to-substrate ratio, 1:20)-mediated proteolysis using the same cycling scheme for 90 cycles. The resultant peptides were cleaned using SEP-PAC C18 (Waters Corp., Milford, MA) and analyzed, after spike-in 10% iRT peptides ^51^, using SWATH-MS following the 32-fixed-size-window scheme as described previously ^19, 21^ using a 5600 TripleTOF mass spectrometer (Sciex) and a 1D+ Nano LC system (Eksigent, Dublin, CA). The LC gradient was formulated with buffer A (2% acetonitrile and 0.1% formic acid in HPLC water) and buffer B (2% water and 0.1% formic acid in acetonitrile) through an analytical column (75 μm × 20 cm) and a fused silica PicoTip emitter (New Objective, Woburn, MA, USA) with 3-μm 200 Å Magic C18 AQ resin (Michrom BioResources, Auburn, CA, USA). Peptide samples were separated with a linear gradient of 2% to 35% buffer B over 120 min at a flow rate of 0.3 μl min^−1^. Ion accumulation time for MS1 and MS2 was set at 100 ms, leading to a total cycle time of 3.3 s.

### SWATH assay query library for prostate tissue proteome

To build a comprehensive library for SWATH data analysis, we analyzed unfractionated prostate tissue digests prepared by the PCT method using Data Dependent Acquisition (DDA) mode in a tripleTOF mass spectrometer over a gradient of 2 hours as described previously ^19^. We spiked iRT peptides ^51^ into each sample to enable retention time calibration among different samples. We then combined these data with the DDA files from the pan-human library project [104]. All together we analyzed 422 DDA files using X!Tandem ^52^ and OMSSA ^53^ against three protein sequence databases downloaded on Oct 21, 2016 from UniProt, including the SwissProt database of curated protein sequences (n=20,160), the splicing variant database (n=21,970), and the trembl database (n=135,369). Using each database, we built target-decoy protein sequence database by reversing the target protein sequences. We allowed maximal two missed cleavages for fully tryptic peptides, and 50 p.p.m. for peptide precursor mass error, and 0.1 Da for peptide fragment mass error. Static modification included carbamidomethyl at cysteine, while variable modification included oxidation at methionine. Search results from X!Tandem and OMSSA were further analyzed through Trans-Proteomic Pipeline (TPP, version 4.6.0) ^54^ using PeptideProphet and iProphet, followed by SWATH assay library building procedures as detailed previously ^19, 55^. Altogether, we identified 167,402 peptide precursors, from which we selected the proteins detected in prostate tissue samples, and built a sample-specific library. SWATH wiff files were converted into mzXML files using ProteoWizard ^56^ msconvert v.3.0.3316, and then mzML files using OpenMS ^57^ tool FileConverter. OpenSWATH[105] was performed using the tool OpenSWATHWorkflow with input files including the mzXML file, the TraML library file, and TraML file for iRT peptides.

### Peptide quantification using OpenSWATH

To obtain consistent quantification of the SWATH files, we obtained the all annotated *b* and *y* fragments from the sp, sv and tr libraries. About ten thousand redundant and low-quality assays were removed. Then we extracted the chromatography of these fragments and MS1 signals using OpenSWATHWorkflow, followed by curation using DIA-expert[106]. Briefly, the chromatography of all fragments and MS1 signals were subject to scrutiny by empirically developed expert rules. A reference sample with best q value by pyprophet was picked up to refined fragments. The peptide precursors are further filtered based on the following criteria: i) remove peptide precursors with a q value higher than 1.7783e-06 to achieve a false discovery rate of 0.00977 at peptide level using SWATH2stats [107]; ii) peptides with a FC higher than 2 between the reference sample and its technical replicate were removed; iii) peptides matching to multiple SwissProt protein sequences were removed. The data matrix was first quantile normalized, log_2_ transformed, followed by batch correction using the ComBat R package [108]. Finally, for each protein and pair of technical replicates the average value was computed.

### Statistical analysis

All plots were produced with R. Kaplan-Meier estimators were used for RFS analysis. Differences between survival estimates were evaluated by the log-rank test.

### Computation of molecular perturbation scores

On the genomic level (mutation and CNA), we kept the tumor samples (66 in total) that contain FCs with respect to the blood. The mutation matrix was further discretized by setting all non-zero events to 1. At the transcriptomics level, the FCs for the 63 tumor samples were computed as described above (see ‘RNA Sequencing’). Finally, on the proteomics level, we computed the FCs for the tumor samples (66 in total) as follows: for each protein, its mean intensity over the normal samples was subtracted from the intensities of the tumor samples. (We chose to compute the FCs for the tumor samples with respect to a global reference (average of all normal samples) and not with respect to their paired benign sample in order to achieve a higher consistency with the transcriptomics level.)

We assigned to each sample two molecular perturbation scores summarizing/quantifying the magnitude of its FCs: DE_count counts the number of mutated/differentially expressed (DE) genes, while the DE_sum score is the sum of absolute FCs of all genes. Thus, while the first score counts the number of events (mutations/DE genes), the second one quantifies their magnitude. These two scores can be regarded as generalizations of the term ‘mutational burden’ for the mRNA and protein layer. A gene is regarded as mutated/DE if its value is 1 in the mutation layer and if its absolute value is above a threshold that has been set to 1 for the mRNA and protein layer. For the CNA layer, the corresponding threshold was set to 0.5 because the range of FCs in the CNA matrix is smaller than the mRNA and protein matrices. Both types of scores were computed for each molecular level, except for the point mutations where only DE_count was computed. Afterwards, the 66 DE_count scores (63 for the mRNA) and the DE_sum scores at each layer were divided into the four grade groups G1, G2, G3 and G4/5 respectively.

### Correlating CNAs with mRNA and protein layer

For each of the 2,120 genes measured in all three layers (CNA, mRNA and protein), we computed the Spearman correlation between its CNAs and corresponding mRNA FCs as well as between its CNAs and corresponding protein FCs. We reduced each layer to the 63 tumor samples with available mRNA data.

### Network propagation/smoothing

As a network, the STRING gene interaction network (version 10)[43] was used, after removing all edges with combined score smaller or equal to 0.9 and keeping subsequently the largest connected component. The resulting network consisted of 10,729 nodes and 118,647 (high-confidence) edges. For the network smoothing, the weight matrix was computed as described in Vanunu et al.[49], but for an unweighted graph and the propagation parameter was set to 0.5. The propagation was iteratively repeated 500 times to ensure convergence of the results. For the mapping from gene symbols to STRING identifiers (**Additional file 7: Table S6**) we used the R/Bioconductor package STRINGdb [109]. The gene symbols with no matching STRING identifier were removed, while for those that mapped to multiple STRING identifiers, the first mapping was kept (default choice in the package). From the multiple gene symbols that mapped to the same STRING identifier, the first mapping was kept. The genes that were not present in the network were removed from the datasets, while those that were present in the network but not in the corresponding dataset were initially filled in with 0’s.

Genes with very small, ‘smoothed’ (absolute) FCs were filtered out as follows: after the network propagation, only network nodes that had protein measurements themselves or at least one direct neighbor (on the filtered STRING network) with protein measurements were considered in the next steps of this analysis. *I*.*e*. network nodes without measured FCs at the protein layer that had no direct neighbor with measured protein values were removed from the subsequent analyses.

For significance testing, the one-sided Wilcoxon rank sum test comparing the smoothed FCs between the groups G4/5 (consisting of 12 samples for the CNA, mRNA and protein layer) and G1 (consisting of 26 samples for the CNA and proteins and 25 for the mRNA) was applied to each network node (after filtering) and layer, once for up-regulation and once for down-regulation. The resulting sub-networks (up-regulated and down-regulated) consisted of those genes that were significant (*P* value below 0.05) at all three layers and all of the edges connecting them on the filtered STRING network.

It should be noted that although measurements from the same patient might not be statistically independent, we have kept them in our analyses firstly in order to increase statistical power and secondly because not all of them correspond to clonal events as shown on **Fig. 5**. To make sure though that having two samples for some of the patients has not affected our conclusions, we have repeated the statistical testing step (one-sided Wilcoxon rank sum test) in two ways: comparing G4/5 with G1 as before but removing the second tumor area of a patient if it belonged to the same grade group as tumor area 1 (*i*.*e*. removing TA2 of patients H3 and H10), and secondly comparing G4/5 with the combined (G1 and G2) group and once again removing the second tumor area of a patient if it belonged to the same grade group as his tumor area 1. *P* values resulting from these analyses were highly correlated (**Additional file 1: Fig. S9**), and we would thus consider the current conclusions to be robust.

### Network Component 1 analysis

For each tumor sample at the CNA layer, a one-sided, one-sample t-test has been applied testing if its average FC over the genes of the Network Component 1 (and in particular those that have been measured at the CNA) is significantly greater than 0. Due to the presence of outliers in some samples, the non-parametric, one-sided Wilcoxon signed-rank test has been applied as well yielding very similar results (data not shown). A result is considered to be significant if the corresponding *P* value is below 0.05. The analysis has been repeated for the mRNA and protein layer.

### Independent cohorts validation

For the validation of Network Component 1, we used published datasets of three PCa cohorts: TCGA, MSKCC, and Aarhus. For TCGA and MSKCC, we downloaded the CNA, mRNA with precomputed z-scores per gene, and corresponding clinical data from cBioPortal[110] (https://www.cbioportal.org/). There were 489 samples with log_2_CNA data and 493 samples with mRNA profiles in TCGA. In MSKCC, there were 157 primary tumors with CNA data and 131 primary tumors with mRNA data. The clinical endpoint used in TCGA was the progression-free survival time and the disease-free survival in MSKCC. All previous samples had known survival time.

For the Aarhus study (NCBI GEO dataset GSE46602), we downloaded the mRNA matrix and corresponding clinical information as described in Ycart et al [111]. The resulting mRNA matrix consisted of 20,186 genes and 50 samples-36 PCa samples with known RFS time and 14 benign samples. Once excluding the benign samples, we computed z-scores per gene in order to have comparable values with the other two studies. These 36 PCa samples were also considered in the subsequent survival analysis. CNA data was not available for the Aarhus study.

We reduced all datasets to the nine genes of Network Component 1. In each of the datasets, we computed for each sample an average copy number change (CNA) or an average z-score (mRNA) across the nine genes of Network Component 1 (combined risk score). Subsequently, we used these combined risk scores to split the samples of each dataset into two groups: samples with a combined risk score larger or equal to the median combined risk score of the study were considered as ‘altered’ and the rest as ‘unaltered’. Kaplan-Meier curves were generated for the two groups. Due to the high level of discretized values in MSKCC at the CNA layer, a sample is considered to be ‘altered’ in that dataset if its combined risk score is above zero.

Additionally, we fitted for each dataset a Cox proportional-hazards model to predict survival time using as input variables the average copy number change (CNA) or average z-score (mRNA) of Network Component 1 (variable of interest) and the age (when available, *i*.*e*. for TCGA and Aarhus). For each dataset with available copy number information (*i*.*e*. for the TCGA and MSKCC studies), we fitted a second Cox proportional-hazards model with the fraction of genome altered as an additional input variable. For the model fitting, we used the R package survival (https://cran.r-project.org/web/packages/survival/index.html).

### Analysis of regulators and target genes

For this analysis, we used once again the STRING gene interaction network. For each target gene (AGR2, ACPP, POSTN, LGALS3BP), we split the network nodes into two groups as follows: firstly we identified the neighbors of the target gene supported by a combined evidence larger than 0.2. This set together with the target gene constituted group 1 while the remaining network nodes constituted group 2. For this splitting, only genes present in the network with copy number measurements and with a matching STRING identifier (**Additional file 7: Table S6**) were considered (*i*.*e*. 17,306 genes in total). Subsequently, genes altered (*i*.*e*. with log_2_ copy number ratio greater than 0.5 in absolute) in fewer than four tumor samples across the 66 tumor samples were filtered out in each of the two groups. Genes in group 1 after the CNA filtering are potential regulators of the target under consideration. For each gene in the two groups after the filtering, we computed the Spearman correlation between its CNAs and the mRNA FCs of the target gene. For computing the correlation, the samples were reduced to the 63 tumor samples with available mRNA data.

Subsequently, we fitted an elastic net model with alpha=0.5 for ACPP. We used as output variable the mRNA FC of ACPP and as input variables the CNAs of the genes in group 1 after the copy number event filtering. The value for the regularization parameter lambda was chosen through 10-fold cross validation (default in the R package glmnet (https://cran.r-project.org/web/packages/glmnet/)). The samples were necessarily reduced to the 63 mRNA tumor samples. Predictors/regulators with a non-zero beta coefficient were deemed significant. We have used the elastic net model with alpha=0.5 because it is a method giving sparse solutions and can deal with correlated predictors at the same time.

As an additional validation to our approach, we used the two independent PCa cohorts described above (TCGA and MSKCC) and reduced the samples to those having both CNA and mRNA profile. This resulted in 488 samples for TCGA and 109 samples for MSKCC. Next, for each of the significant regulators/predictors we computed the Spearman correlation between its CNAs and the corresponding mRNA z-scores of ACPP in each of the two independent studies and checked if the sign of the Spearman correlation matched the sign of the Spearman correlation computed for our cohort, *i*.*e*. there was an agreement regarding the direction of the association.

## Declarations

### Ethics approval and consent to participate

This study was approved by the Cantonal Ethics Committee of Zurich (KEK-ZH-No. 2008-0040), the associated methods were carried out in accordance with the approved guidelines, and each patient has signed an informed consent form.

### Consent for publication

Not applicable.

### Availability of data and materials

Exome and RNA sequencing data were submitted to the Sequence Read Archive (SRA) at NCBI under accession numbers PRJNA577801 (exome-seq) and PRJNA579899 (RNA-seq), respectively. The SWATH proteomics data were deposited in PRIDE. Project accession code is PXD004589. The published datasets of the two PCa cohorts (TCGA and MSKCC) analyzed during the current study can be downloaded from cBioPortal[110] while the third (Aarhus) is available at the NCBI GEO repository under the accession number GSE46602.

### Authors’ contributions

A.B., T.G., P.J.W. and R.A. designed the project. P.J.W., T.G., Q.Z., C.E.F., N.J.R., A.C, D.R, J.H.R., C.F., K.S., C.P., T.H., A.L.M. and C.B. procured the samples and performed the experiments. K.C., T.G., Q.Z., U.W., R.S., N.C.T, K.O., L.C., L.M., M.R.M, M.M and A.B. designed and performed the statistical analyses with critical inputs from C.Y., H.C., Q.Z., Y.Z., M.H. and other authors. K.C., A.B., T.G. and R.A. interpreted the results. K.C., T.G., P.J.W., A.B. and R.A. wrote the manuscript with inputs from all co-authors. A.B., R.A., P.J.W. and T.G. supported and supervised the project.

## Acknowledgements

Not applicable.

## Supplementary information

**Additional file 1: Supplementary text and supplementary figures**.

**Additional file 2: Table S1. Clinicopathological, immunological and other molecular information of the 39 PCa patients**. (a) Overall clinicopathological characteristics. (b) Detailed information for each patient. Pat: numeric patient ID; Pat_id: patient ID grouped by the overall grade. L: low grade; M: intermediate grade; H: high grade; Overall_Gleason_GrGp: overall ISUP grade group; pT: tumor stage; pN: nodal status; R: surgical margin status; Age_at_OP: age at operation; PSA_at_Diag: blood PSA level at diagnosis; Time (months): RFS time. A value of 0 corresponds to patients excluded for the reasons explained in the ‘**Methods**’ section (see ‘Patients and samples’); Status: status indicator. 1 means recurrence; DX name: tissue region name; ImageName: name of the scanned images; index_tumor_id: patient ID of TA1 (or TA); TA1_GrGp: grade group for TA1; T_GrGp: grade group for TA2.

**Additional file 3: Table S2. Exome analysis of the peripheral blood cells and 105 prostate tumor punches in 39 patients**. (a) Allele frequencies (AF) of somatic single nucleotide variants (SNVs) that were called by our bioinformatics pipeline. Genes with called SNV are indicated by an AF > 0. A value of 0 indicates that no SNV was found in the respective genes. In our data, no gene was found with more than one called somatic SNV. (b) Number of samples per gene with called somatic SNV. (c) Protein domain analysis using DAVID.

**Additional file 4: Table S3. Copy number analysis of 105 PCa samples**. (a) Log_2_ ratios indicating the CNA status are shown for all genes in all samples. Values were determined by overlapping gene locations with CNA segments as calculated by CopywriteR. In case more than one segment overlapped with a gene, number was chosen that had the highest absolute value. (b) Genes are shown with log_2_ ratios higher than 0.5 or lower than -0.5 in at least one sample.

**Additional file 5: Table S4. RNA-seq analysis**. (a) Log_2_FCs (relative to all benign samples) for all genes across the tumor samples. (b) mRIN score per sample generated using mRIN (v1.2.0). (c) ETS family gene fusions observed in tumor samples using FusionCatcher: a value of 1 means that the fusion was observed in the respective sample but not its corresponding benign sample, otherwise the value is 0. (d) Normalized RNA-seq count data matrix.

**Additional file 6: Table S5. Proteomics data of 210 PCa samples with duplicates**. (a) Sample information includes patient ID, clinical diagnosis, sample ID and batch design. (b) Protein matrix of log_2_ scaled intensity of 2,371 proteins quantified in 210 PCa samples.

**Additional file 7: Table S6. Integration analysis of 66 tumor samples**. (a) Information (i.e. reference linking them to PCa, consistency between observed and reported effect and number of tumor samples with CNAs) for the first 10 highest-scoring proteins (those with largest average absolute FCs across all tumor specimens). (b) Consistently up-regulated genes in the high-grade tumors: for each of these genes, there is a significant up-regulation of its FCs after network smoothing in the group G4/5 compared to the group G1 in all three layers (CNA, mRNA and protein). (c) Consistently down-regulated genes in the high-grade tumors: for each of these genes, there is a significant down-regulation of its FCs after network smoothing in the group G4/5 compared to the group G1 in all three layers (CNA, mRNA and protein). (d) Chromosome information for the gene members of Network Component 1. (e) Mapping from gene symbols to STRING identifiers.

