## Supplemental file for "Convergent network effects along the axis of gene expression during prostate cancer progression"

**Supplementary Text**

**Exome and CNA analysis**

Exome data from 105 tissue specimens and 39 peripheral blood samples were obtained by targeted enrichment of exome-coding parts of the genome coupled with next generation sequencing, achieving a mean coverage of 64x per sample with an average of 88% of the bases in the target regions covered with at least 20 reads. Using the GATK data processing pipeline, and the combination of the somatic variant caller programs MuTect and Strelka, we detected relatively few somatic mutations in our tumor samples, which was consistent with a previous report [1]. In total, we detected 988 variants in 806 genes across 105 specimens (**Additional file 3: Table S2**). The mutation pattern across the 105 samples was remarkably sparse (**Additional file 1: Fig. S2**), indicating that for our samples, the mutational background in terms of SNVs was diverse, in agreement with previous reports of independent cohorts [1, 2]. Mutations in SPOP, FOXA1, and MED12 reported in independent cohorts [1, 2] were confirmed in this cohort. We found mutations of ALPK2, DNAH10, FAT4 and HCN1 in four out of 105 samples. Moreover, we found 18 genes, including CDH16 and TP53, mutated in three samples, and 129 genes including CTNNB1, mutated in two samples. The remaining mutations occurred in only one sample. Although the mutation frequency of SPOP (3%) was smaller in our cohort, the mutation frequencies of TP53 (around 5%) and MED12 (3%) were in accordance with published data [1,10].

Based on the exome sequencing data, we detected CNAs throughout all tissue samples using the respective blood samples as reference. The CNA frequency profile of our cohort looked highly similar (**Additional file 1: Fig. S11**) to the respective profiles in published PCa data [4]. The most frequent copy number losses were those of chromosomes 6q, 8p, 10q, 13p, and 16q which is in agreement with the literature [4]. Likewise, according to Williams and colleagues, the most frequent copy number gains in PCa are found in chromosome arms 8q and 3q and the entire chromosome 7, which is in complete accordance with our data (**Additional file 1: Fig. S11**). The CNA number landscapes differed markedly between benign and tumor tissue, agreeing with the literature [3, 4]. By setting a threshold for the absolute log_2_ ratio between sample and reference of 0.5, we discovered that the copy numbers of 6,562 genes were substantially altered in at least one sample (**Additional file 4: Table S3**). 1,110 genes showed copy number gain in at least five samples or copy number loss in at least five samples (**Additional file 1: Fig. S3**). Due to the extensive nature of the CNAs as oftentimes whole chromosome arms are altered, many genes can be categorized into groups of similar CNA patterns. **Additional file 1: Fig. S4** shows the CNA status of signature genes representing known areas of recurrent CNAs in PCa- splitted into fusion-partner and non-fusion-partner genes- for instance loss of PTEN and gain of MYC in high-grade PCa [5].

**RNA-seq analysis**

We measured the transcriptome of all tissue samples using RNA-seq technology. mRNA was extracted from tissue corresponding to the 105 tissue samples described above: 39 normal prostate and 66 tumor tissue samples. 31 out of 105 samples showed strong RNA degradation (**Additional file 1: Fig. S12**). After 3’ bias correction, four samples (BPH, TA1 and TA2 of the patient M5 and TA2 of the patient M8) were excluded from subsequent analyses due to strong remaining bias.

We looked for genes commonly differentially expressed across the tumor samples. We identified a subset of the 31 genes significantly over-expressed in all 63 tumor samples (**Additional file 1: Fig. S5**). Interestingly, ANKRD34B is a patented transcript biomarker for PCa (*Novel rna-biomarkers for diagnosis of prostate cancer*. US 20160298199 A1). PRAME (Preferentially Expressed Antigen in Melanoma) is a tumor-associated antigen involved in the progression of several tumors including PCa [6] and an effective target for immunotherapies [7]. Our data implies PRAME-targeted immunotherapy might be effective for PCa. ADAM7 was reported as a potential biomarker for PCa [8]. Up-regulation of ZIC2 and ZIC5 has also been reported in an independent PCa cohort, while significant correlation of ZIC5 expression with PCa survival has been shown [9]. The most down-regulated transcripts were DAPL1 and ORM2.

We further identified somatic fusions from the RNA-seq data. A large fraction of the tumors harbored ETS family gene fusions (36 out of 63 tumors: 31 x ERG (49.2%), 2 x ETV1 (3.2%), 0 x ETV4, 2 x ETV5 (3.2%), 1 x FLI1 (1.6%)), which are frequently detected in PCa [10]. ETS fusions were mutually exclusive and appeared in tumors from all grade groups (**Additional file 1: Fig. S6**; **Additional file 5: Table S4**).

**Proteome analysis using PCT-SWATH**

We collected tissue punches from 105 tissue regions from the 39 patients and analyzed each punch in technical duplicates using PCT-SWATH [11]. Altogether the 210 samples were processed in 35 batches (**Additional file 6: Table S5**) to eliminate discrepancy in terms of patient identity, tissue type and process date. To interpret the SWATH data, we analyzed the data using OpenSWATH software [12] and a prostate tissue library generated from 422 shotgun analysis of prostate tissue samples, and further refined using DIA-expert [13]. After stringent filtering, we obtained precise quantification for 2,371 SwissProt proteins from 12,957 proteotypic peptides (**Additional file 1: Fig. S13**, **Additional file 6: Table S5**).

**Supplementary Figures**

**Figure S1**

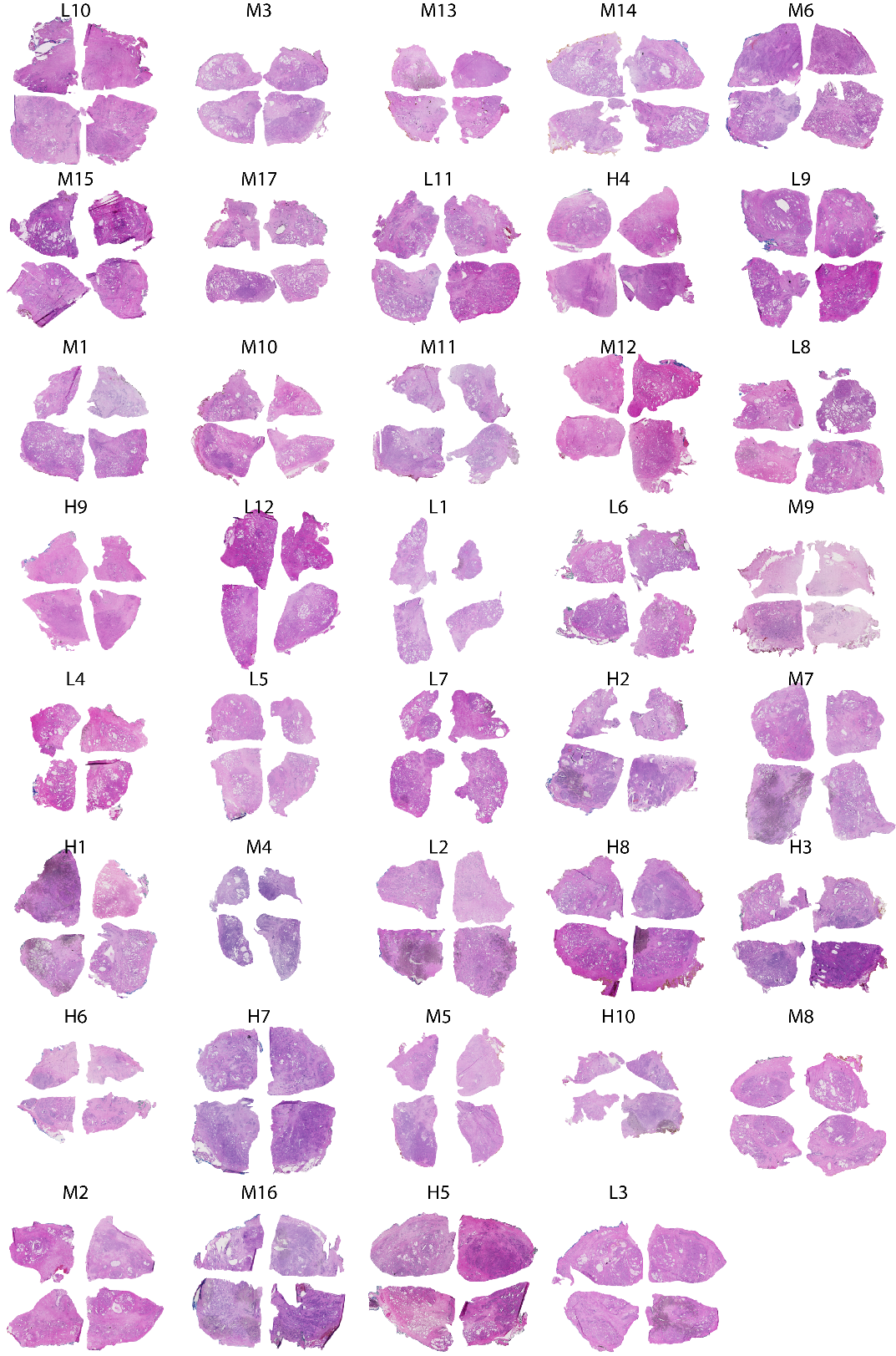

**Figure S1. Experimental design.** H&E staining of the procured fresh frozen prostate tissue of the 39 PCa patients.

**Figure S2**

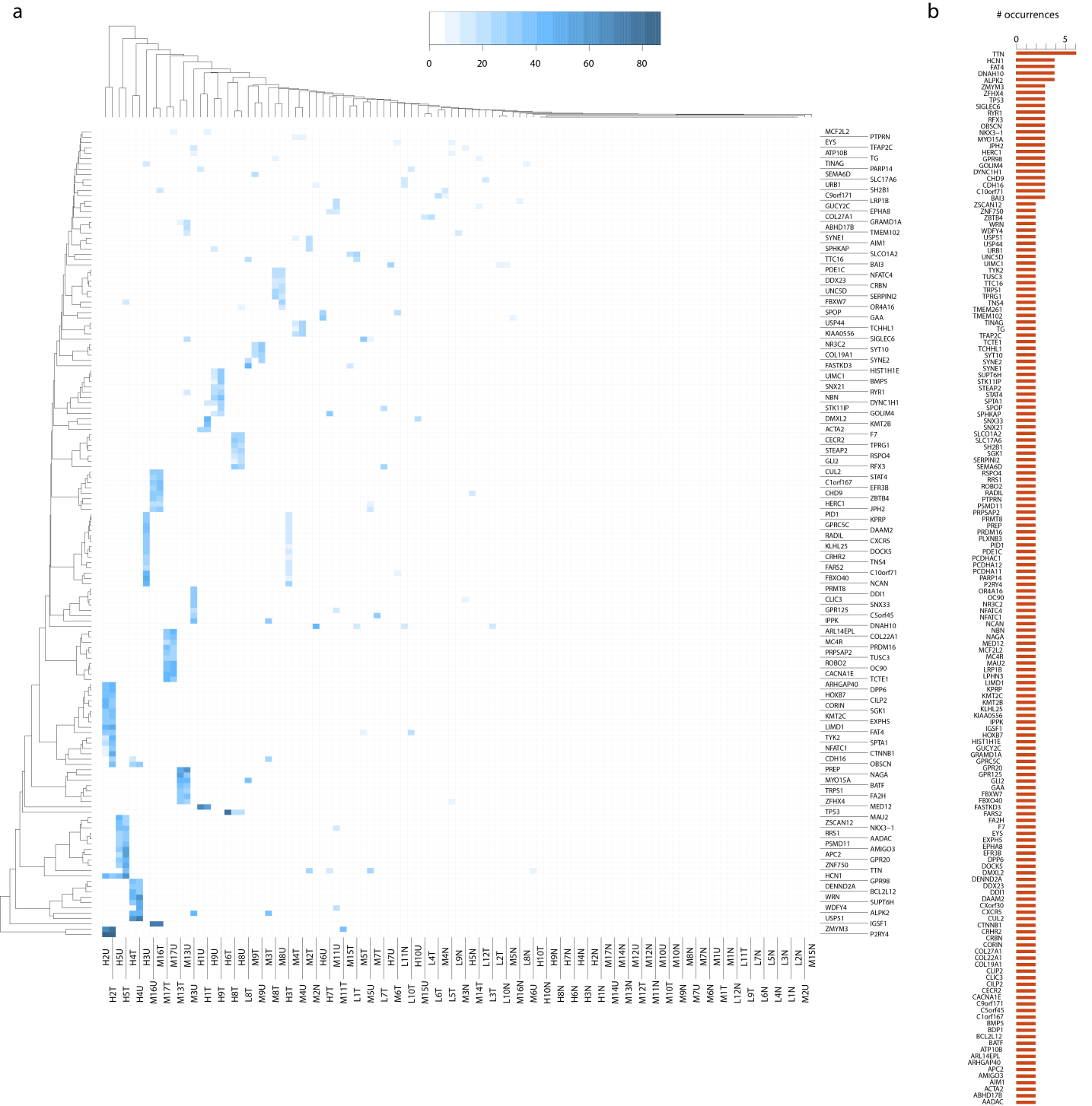

**Figure S2. Heatmap of occurrence of statistically significant point mutations in genes and numbers of occurrence of SNV mutations in individual genes. (a)** Allele frequencies of significant mutations are shown for genes that were mutated in at least two samples within the cohort. In the sample names, the last character N stands for BPH, T for TA or TA1 and U for TA2. The statistical significance was determined by the softwares MuTect and Strelka. **(b)** Bar plot indicating the number of occurrence of SNVs in individual genes throughout 66 tumor (TA, TA1 and TA2) samples in total.

**Figure S3**

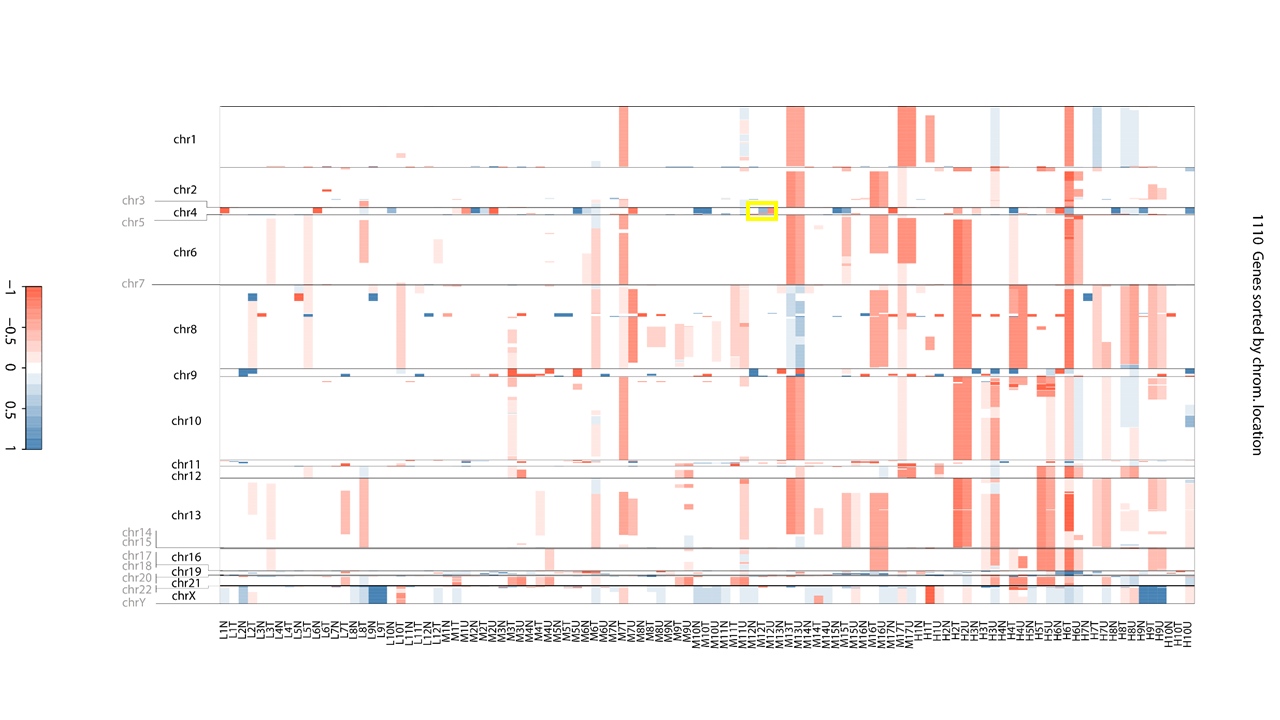

**Figure S3. CNA status of genes with substantial copy number changes.** Heatmap of log_2_-ratio-based copy number changes throughout all samples of 1,110 genes that show copy number loss in at least five samples or copy number gain in at least five samples. Copy number gains are colored blue, copy number losses are colored red. A yellow box has been drawn to mark the ‘interesting’ region for patient M12. For each patient, its representative regions (BPH and TA or BPH, TA1 and TA2) are shown next to each other while patients are ordered based on the grade group (low, intermediate, high). For visualization purposes, all values (*i.e.* log_2_-ratio-based copy number changes) above (below) 1 (-1) have been set to 1 (-1). In the sample names, the last character N stands for BPH, T for TA or TA1 and U for TA2. Chromosomes in gray had too few or no CNA events to be shown.

**Figure S4**

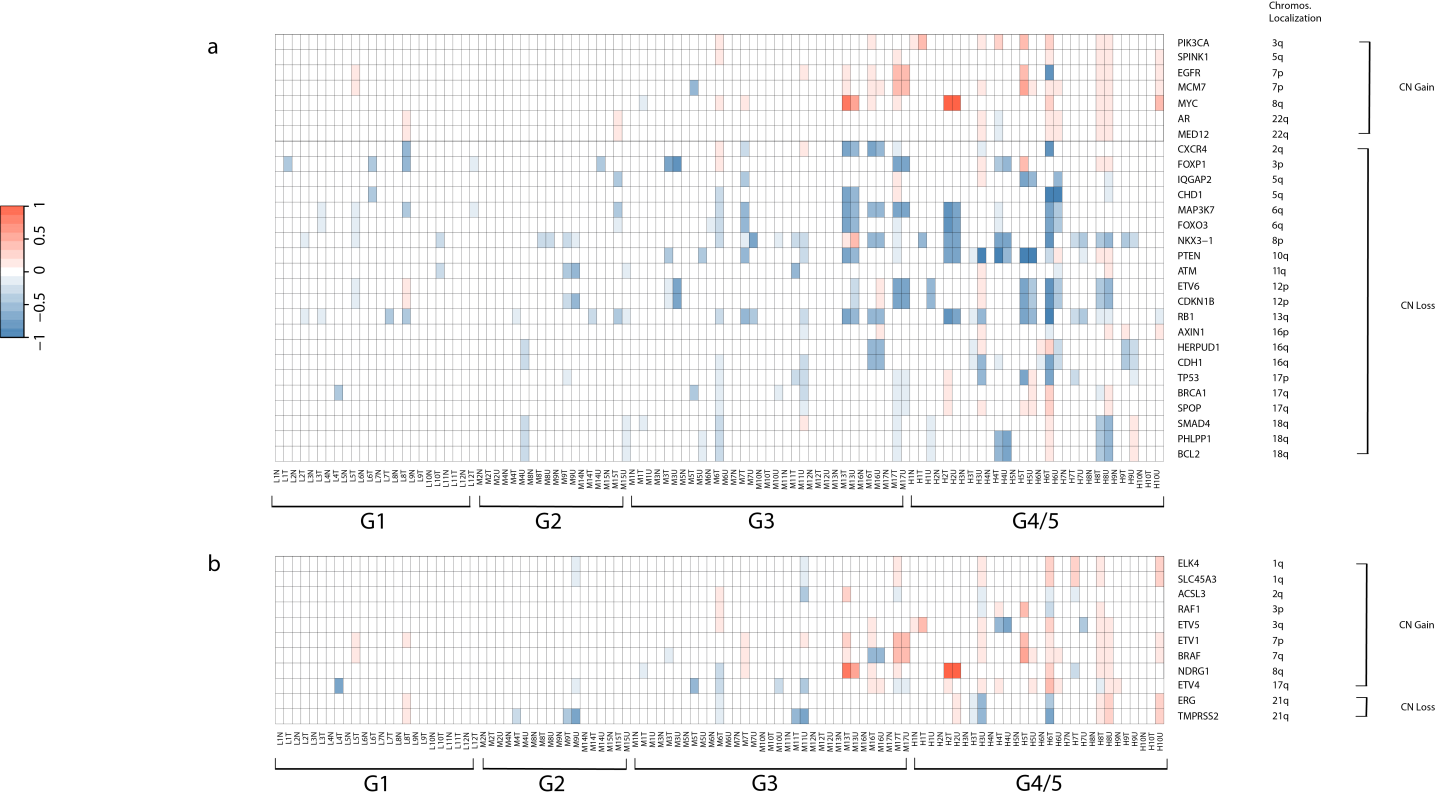

**Figure S4. CNA status of signature genes typically changed in PCa. (a)** Heatmap of log_2_-ratio-based copy number changes throughout all samples for selected genes that are typically changed in PCa samples [4]. Copy number gains are colored red, copy number losses are colored blue. For each patient, its representative regions (BPH and TA or BPH, TA1 and TA2) are shown next to each other while patients are ordered based on the grade group of their higher-grade nodule. For visualization purposes, all values (*i.e.* log_2_-ratio-based copy number changes) above (below) 1 (-1) have been set to 1 (-1). In the sample names, the last character N stands for BPH, T for TA or TA1 and U for TA2. **(b)** Heatmap of log_2_-ratio-based copy number changes throughout all samples for selected potential fusion-partner genes that are typically changed in PCa samples.

**Figure S5**

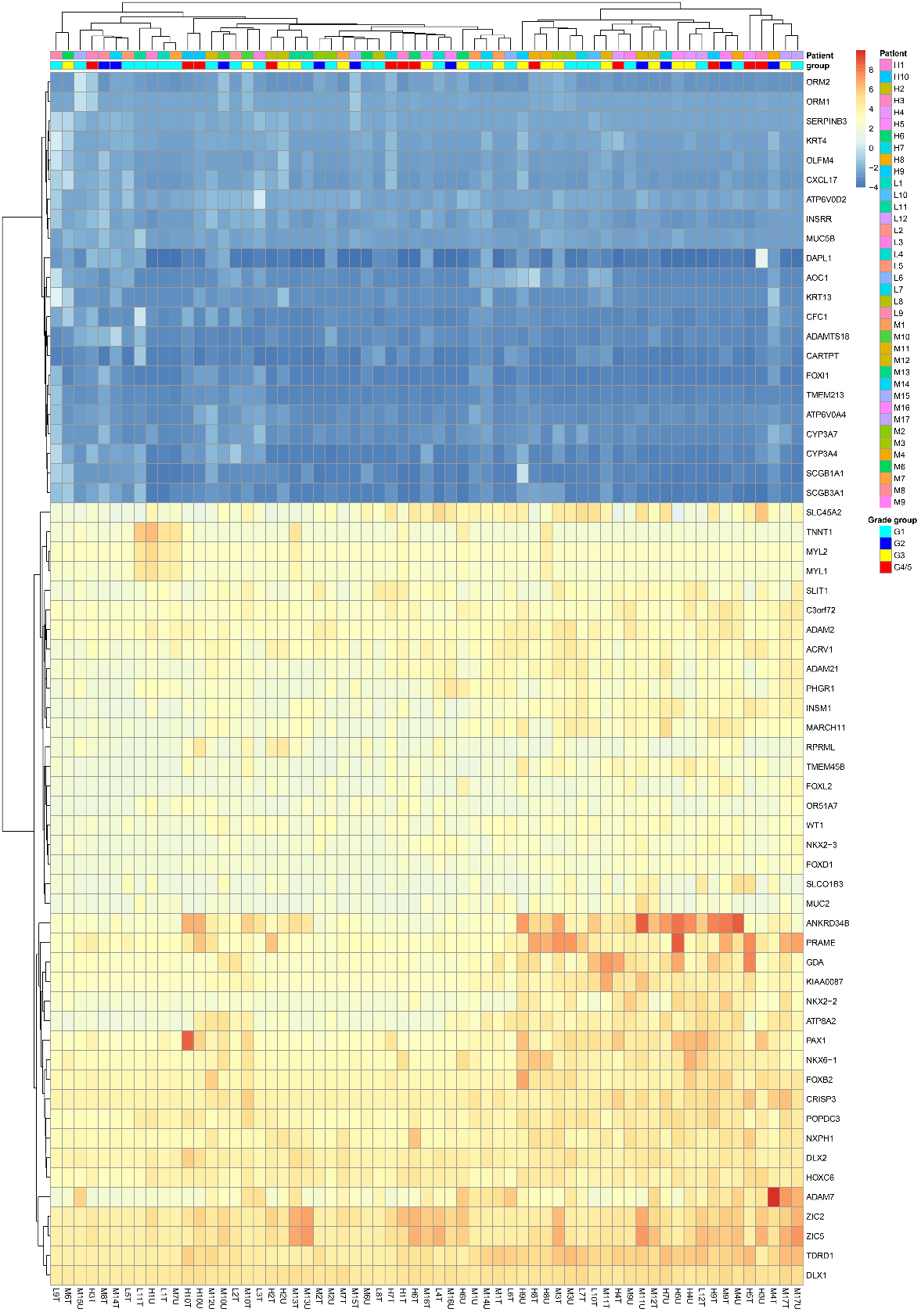

**Figure S5. RNA-seq differential expression analysis.** Log_2_FC (relative to benign tissues) of genes commonly differentially expressed across the tumor samples. In the sample names, the last character T stands for TA or TA1 and U for TA2.

**Figure S6**

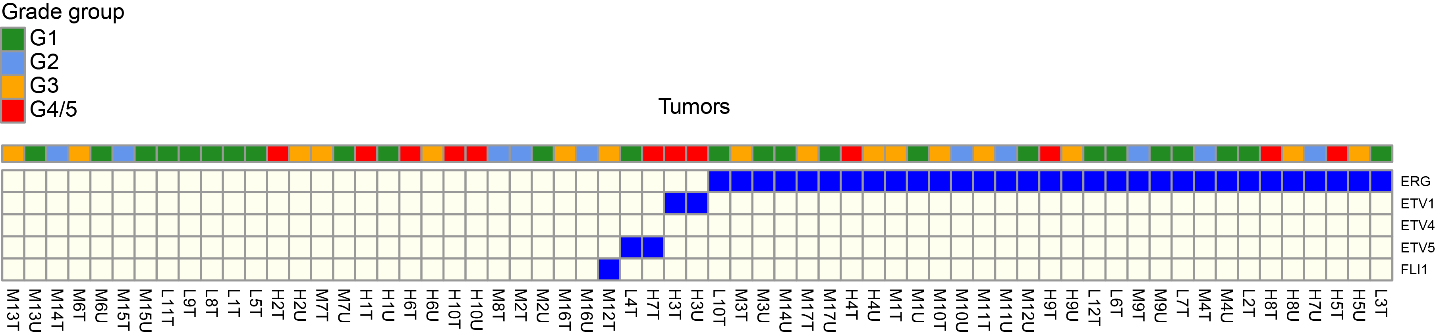

**Figure S6. Gene fusions with ETS genes in tumor samples.** In the sample names, the last character T stands for TA or TA1 and U for TA2.

**Figure S7**

**a**

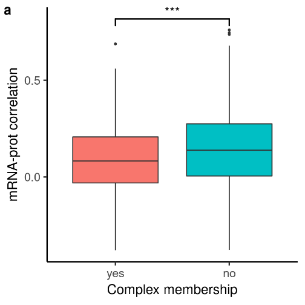

**b**

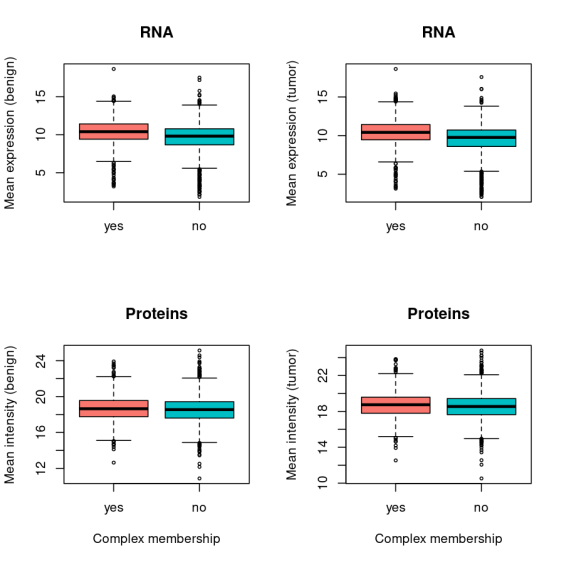

**c**

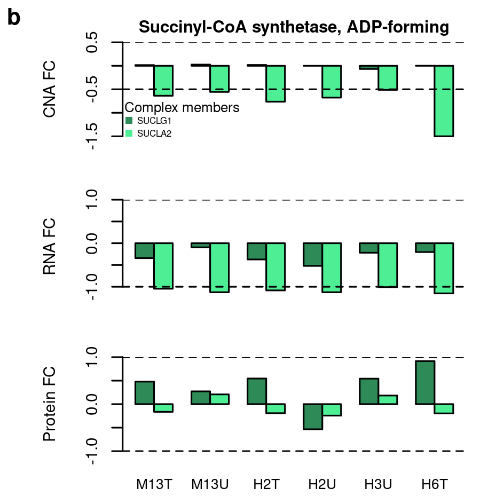

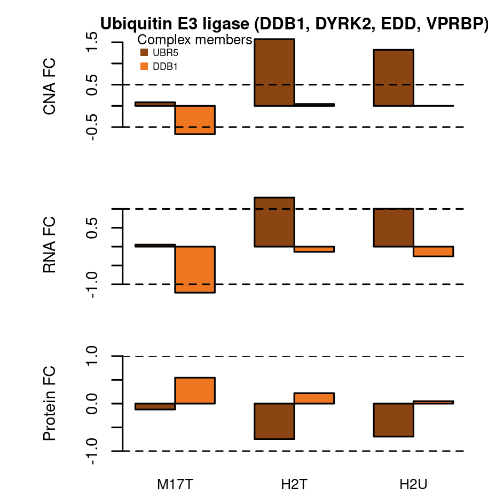

**Figure S7. Protein complexes and buffering.** We classified the measured proteins based on their membership in known protein complexes as ‘inside complexes’ or ‘outside complexes’. The division of the proteins is based on the human ‘core’ protein complexes of the CORUM database[14] (Release July 2017). Out of the 2,371 proteins, there are 789 proteins inside complexes and (the remaining) 1,582 proteins outside complexes. Out of the 789 proteins inside complexes, 724 were also detected on the mRNA layer, while out of the 1,582 proteins outside complexes, 1,399 were detected on the mRNA layer. Correlation analysis was conducted using those 63 tumor samples with available mRNA and protein data. **(a)** Distributions of the mRNA-protein (Pearson) correlations. Pearson correlations of mRNA fold changes versus protein fold changes across all 63 tumor samples for proteins inside (‘yes’) and outside (‘no’) complexes; *** *P* value ≤ 0.001. **(b)** To exclude for the possibility that the stronger correlations are the result of biased mean signal intensities in the two groups, we computed the mean signal (mRNA: top and protein: bottom) of the genes in the two groups. Here, we separated tumor and benign specimens. We observe that the differences between the two groups are very small. In fact, the mRNAs and proteins of complex members have on average slightly stronger signals, which would potentially reduce the technical noise. However, we observe that protein complex members exhibit weaker correlations. Thus, it is unlikely that the different correlations are merely due to variation in technical noise. (c) Two examples of complexes with evidence for buffering (*i.e.* there is regulation of at least one subunit at the CNA and mRNA but no regulation for the proteins): Succinyl-CoA synthetase (subunits: SUCLG1; SUCLA2) (left) and ADP-forming and the cancer related complex Ubiquitin E3 ligase (DDB1, DYRK2, EDD, VPRBP)[15] (subunits: UBR5; DDB1; DYRK2; VPRBP) (right). Bar plots illustrate the FC of the subunits (present in all three layers) for samples where protein buffering was observed, at the CNA (top), mRNA (middle) and protein (bottom) layers. The two dashed lines in each plot have ordinate equal to the threshold for differential expression (±0.5 for the CNA and ±1 for the mRNA and proteins). In the sample names, the last character T stands for TA1 and U for TA2.

**Figure S8**

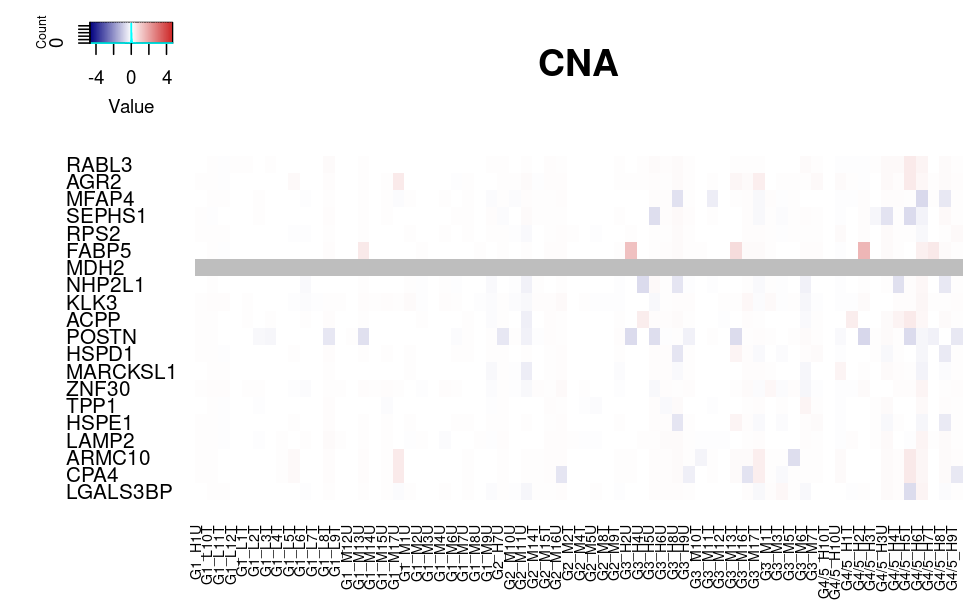

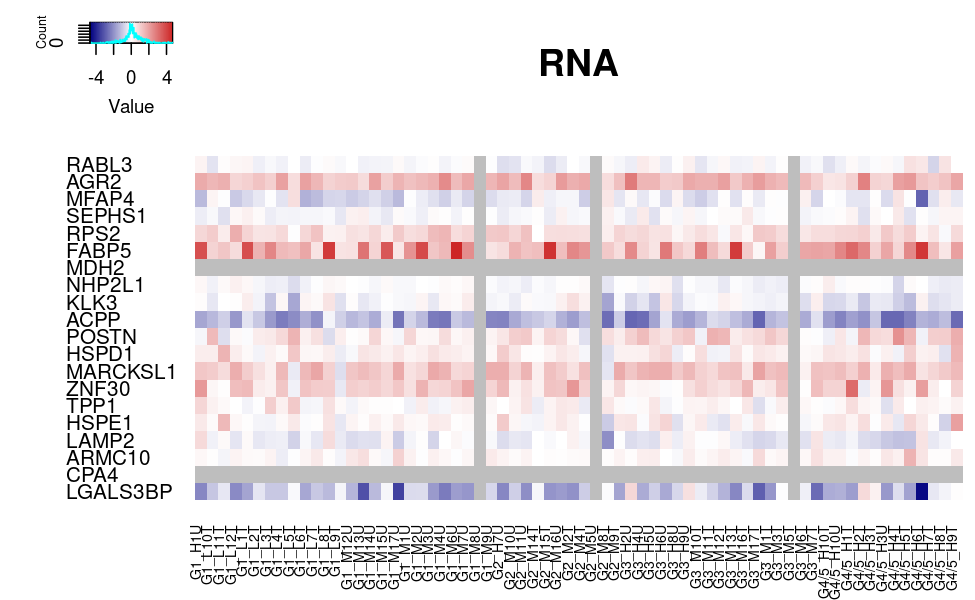

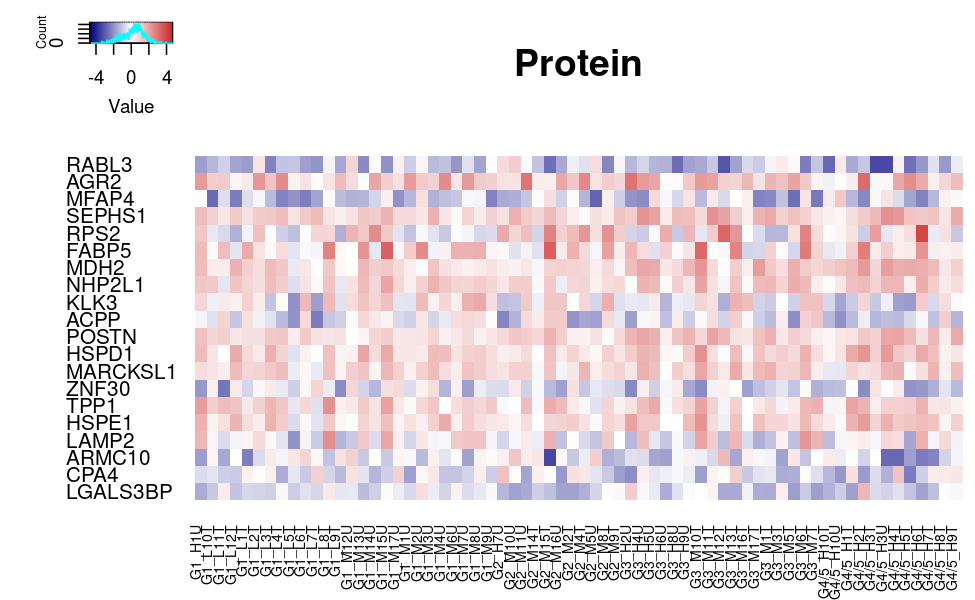

**Figure S8. Heatmaps for the top 20 highest scoring proteins.** Heatmaps of the FCs for the 20 highest scoring proteins across the tumor samples at the CNA, mRNA and protein layer. The columns/samples are ordered based on the grade group while the rows/genes are ranked in decreasing order of the score (score: mean of the absolute protein FCs across all tumor samples). Gray color has been used for genes and samples not being measured at the corresponding layer. In the sample names, the last character T stands for TA or TA1 and U for TA2.

**Figure S9**

**a**

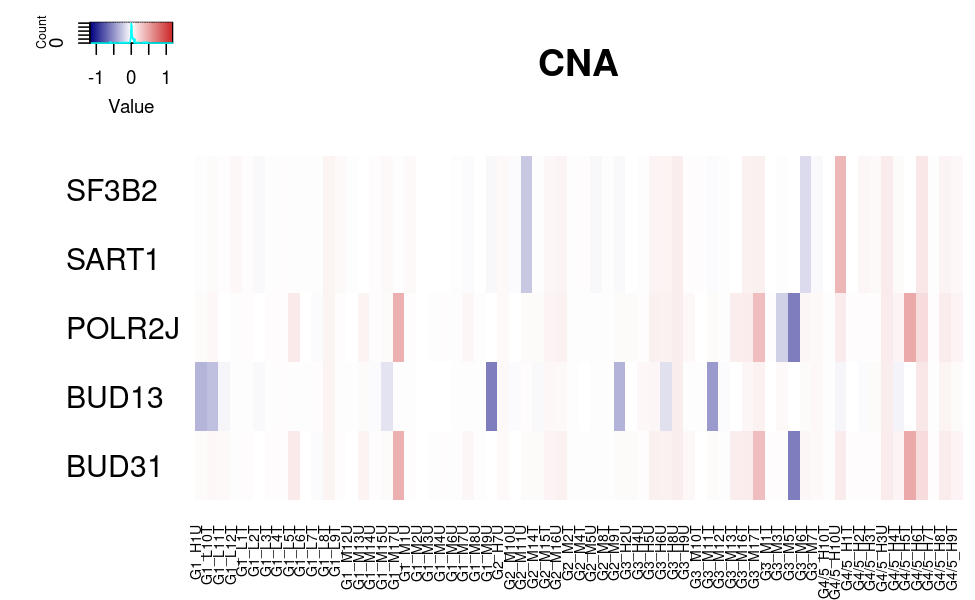

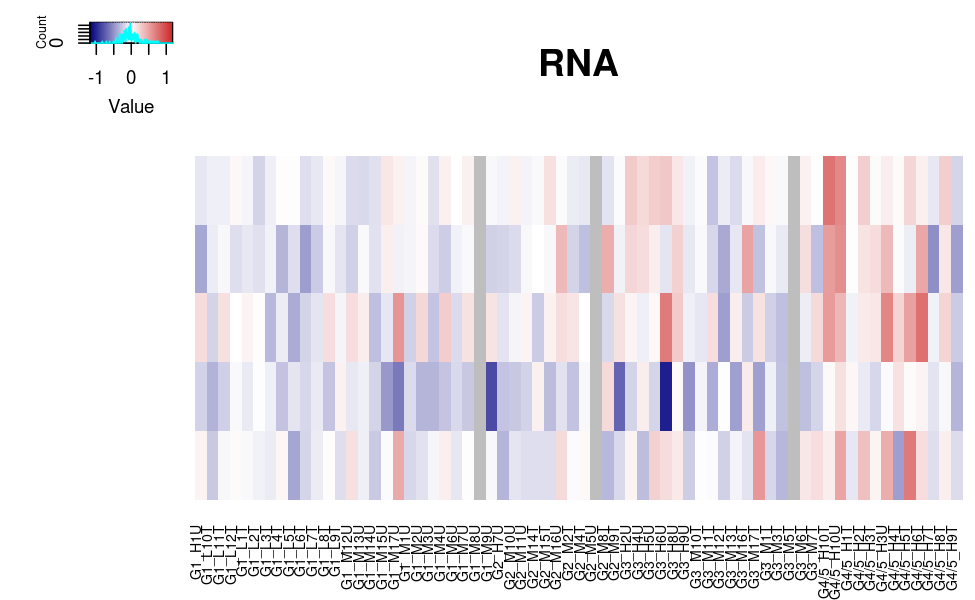

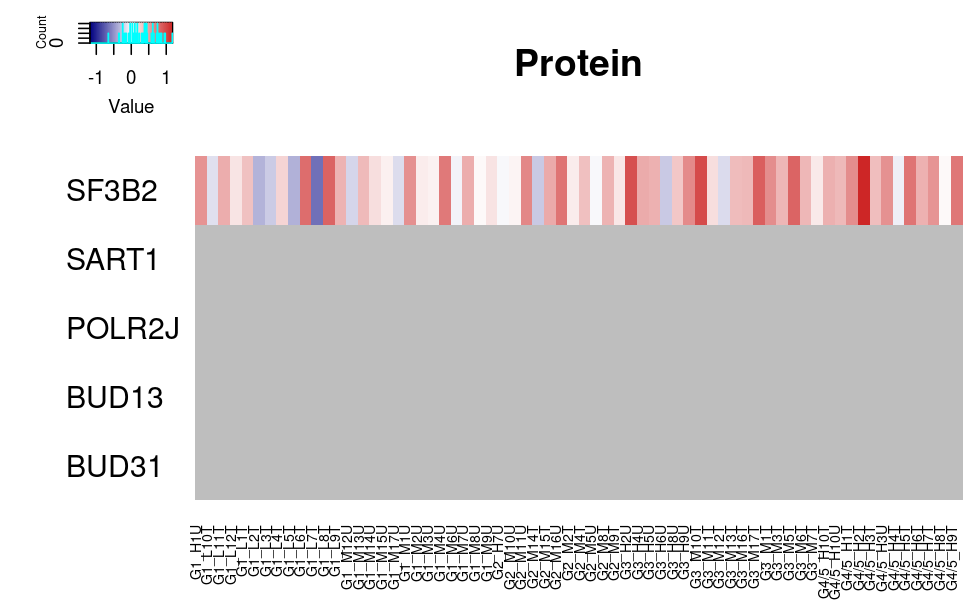

**b**

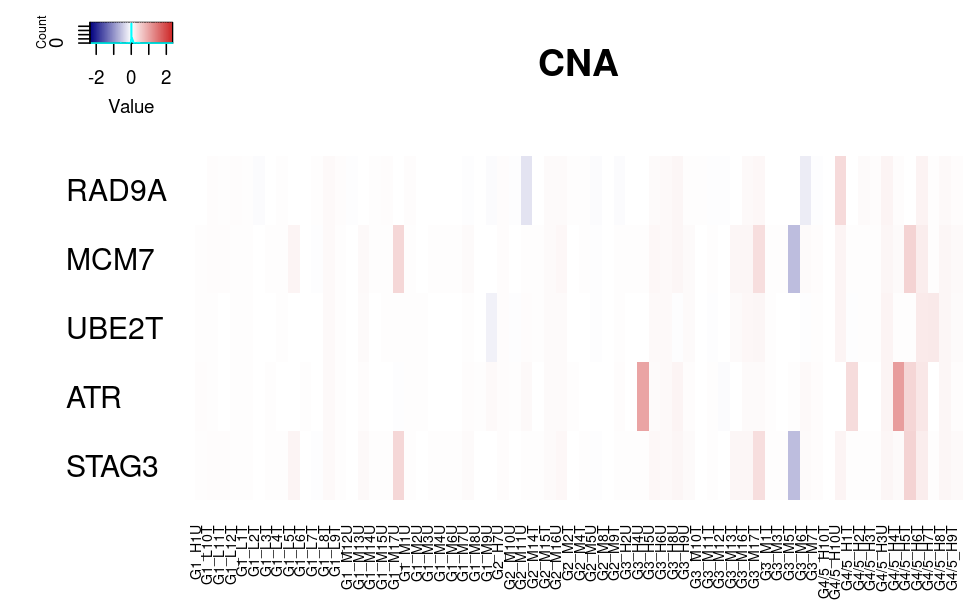

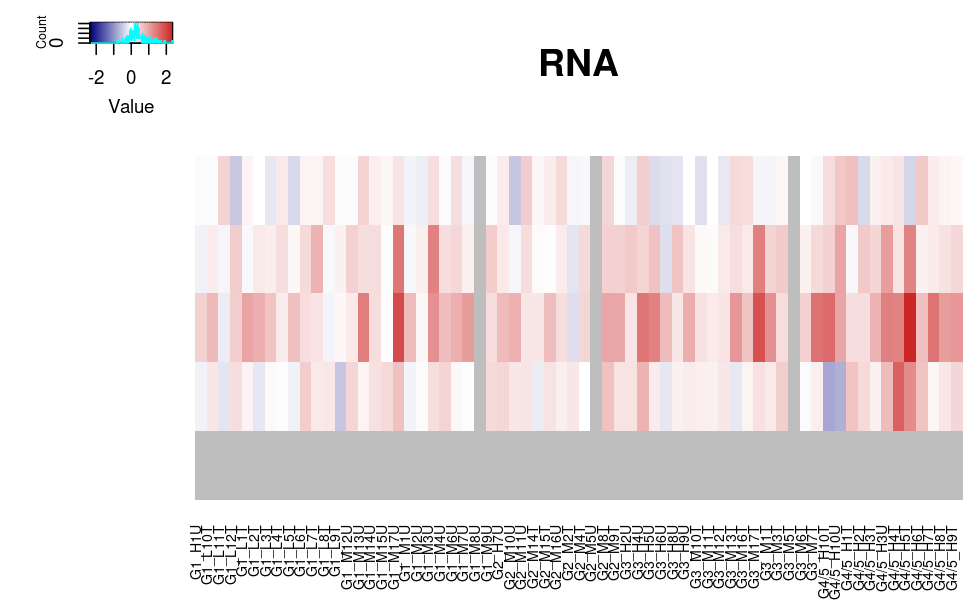

**Figure S9 (*continued*)**

**c	d**

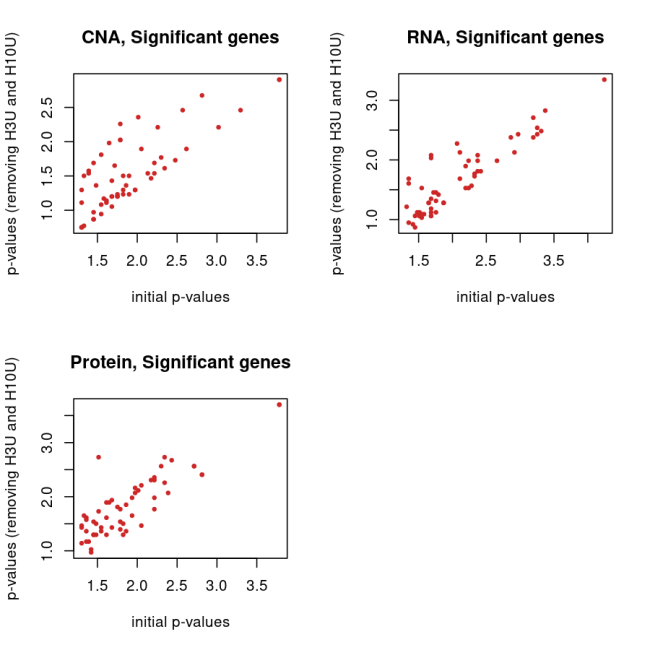

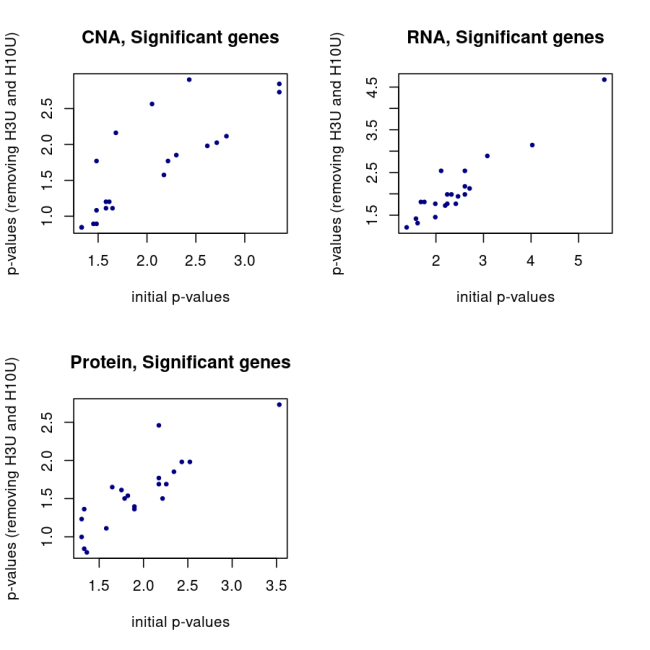

**e	f**

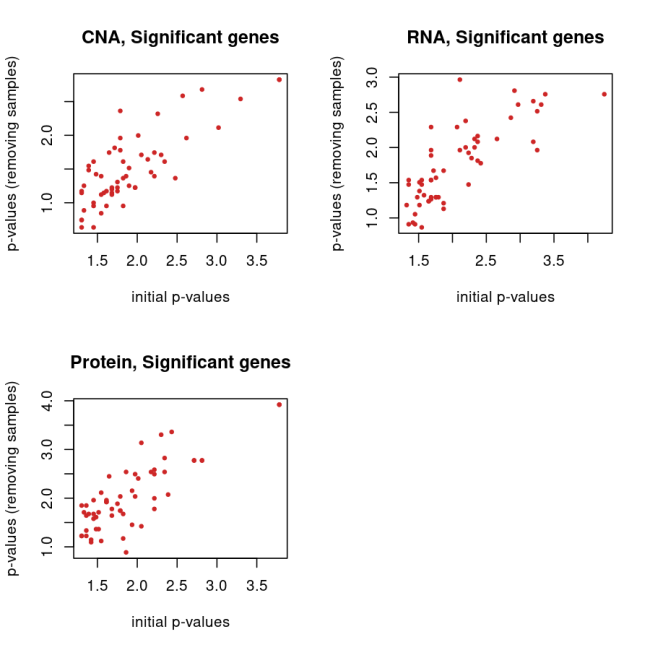

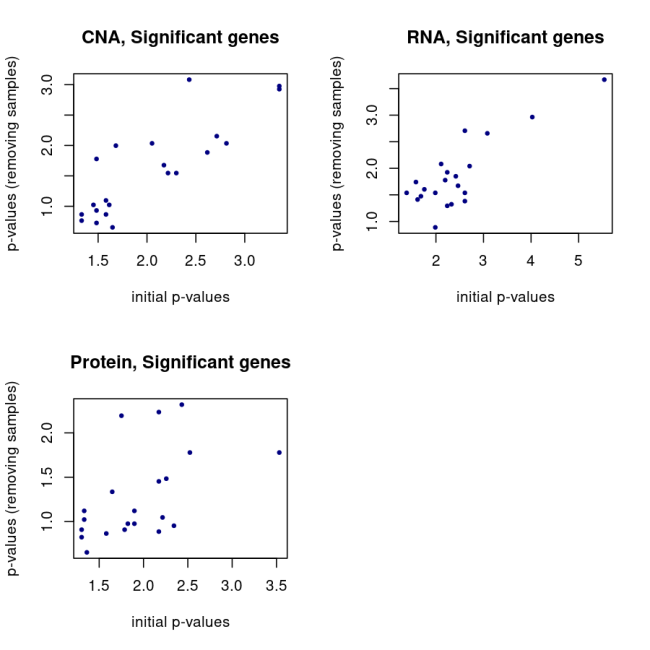

**Figure S9. Heatmaps for selected sub-networks and *P* values from the network smoothing analysis with the two modified approaches.** Heatmaps of the fold-changes (FCs) for Network Component 7 **(a)** and Network Component 8 **(b)** (with up-regulated genes) at the CNA, mRNA and protein layer. The columns/samples are ordered based on the grade group. Only those genes of the sub-network that have been measured in at least one layer are shown. Gray color has been used for genes and samples not being measured at the corresponding layer. If none of the genes of the sub-network has protein measurements, then no protein heatmap is shown. In the sample names, the last character T stands for TA or TA1 and U for TA2. **(c)** Scatterplot between initial *P* values from the network smoothing analysis and corresponding *P* values from the first modified approach (*i.e.* comparing G4/5 with G1 but removing TA2 of patients H3 and H10; **‘Methods’** section)- accounting for potentially dependent measurements- on the CNA, mRNA and protein layer for the genes with up-regulation (upper left). The dots correspond to the significant genes. **(d)** The same is shown on the right for the genes with down-regulation. The *P* values from the second approach (see the **‘Methods’** section for details) are displayed below for up- **(e)** and down-regulation **(f)**.

**Figure S10**

| **a** Group 1 (high similarity at all layers) | | | | | **b** Group 2 (low similarity at all layers) | | |
| --- | --- | --- | --- | --- | --- | --- | --- |
| Patient | Replicate | | TA1 | TA2 |  | TA1 | TA2 |
| H2 | R1 | 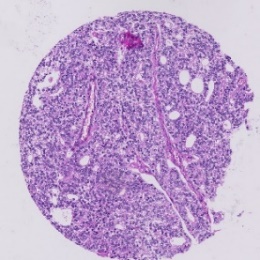 | | 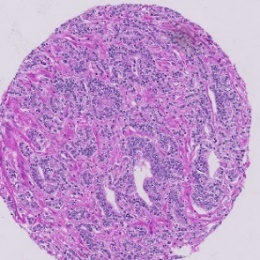 | M12 | 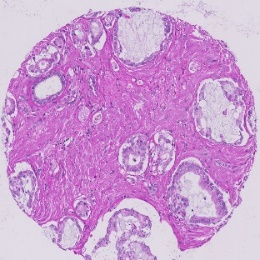 | 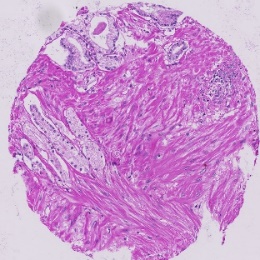 |
|  | R2 | 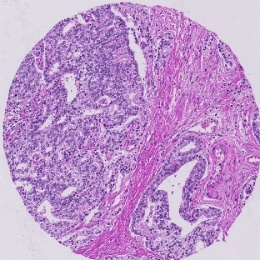 | | 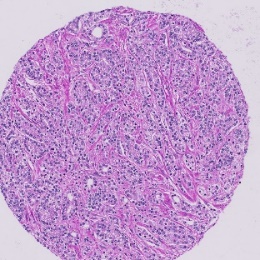 |  | 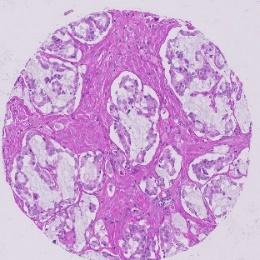 | 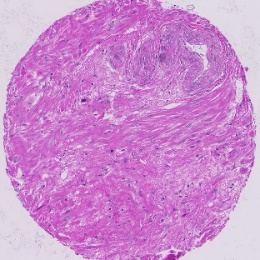 |
| H4 | R1 | 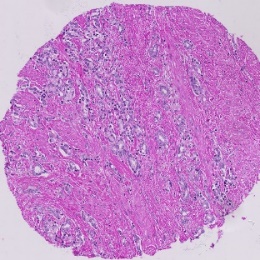 | | 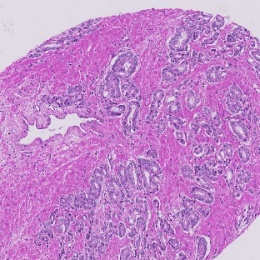 | M14 | 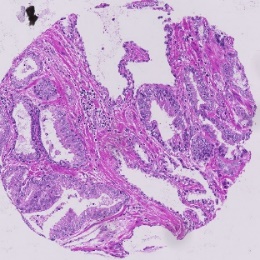 | 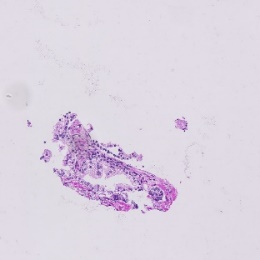 |
|  | R2 | 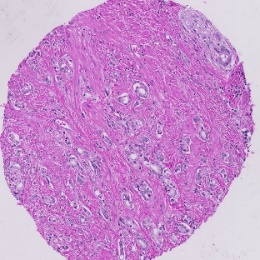 | | 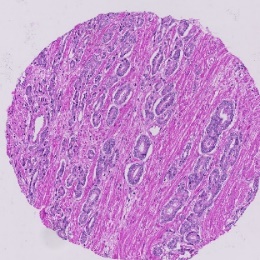 |  | 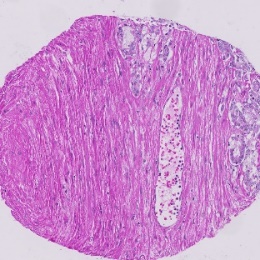 | 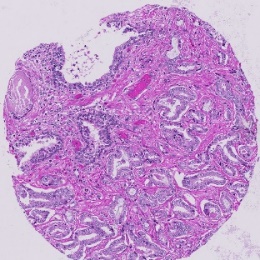 |
| M13 | R1 | 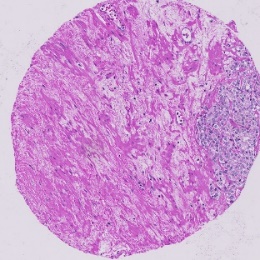 | | 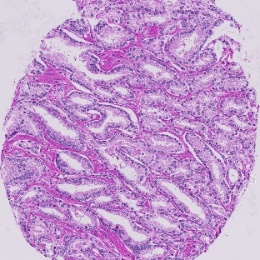 |  |  |  |
|  | R2 | 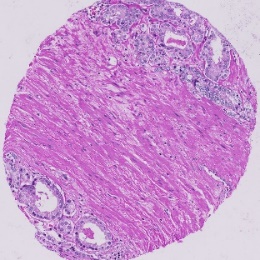 | | 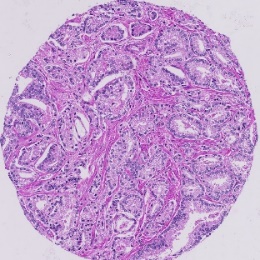 |  |  |  |
| **c** Group 3 (low similarity for the proteins only) | | | |  | **d** Group 4 (high similarity for the proteins only) | | |
| M9 | R1 | 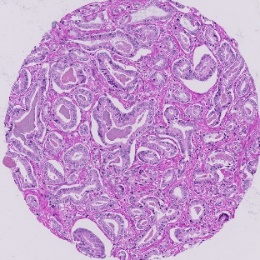 | | 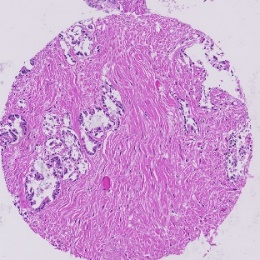 | M6 | 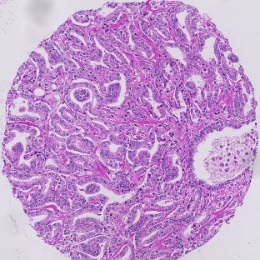 | 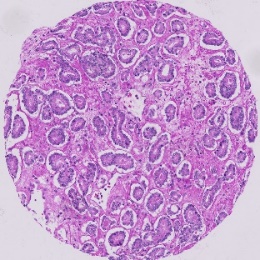 |
|  | R2 | 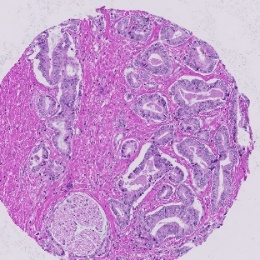 | | 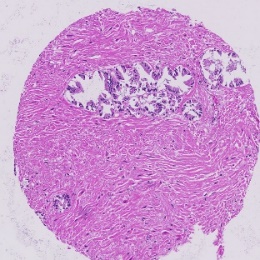 |  | 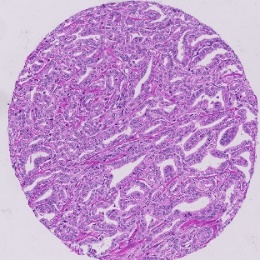 | 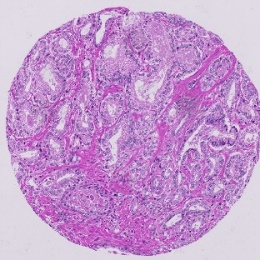 |
| M17 | R1 | 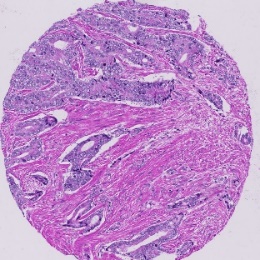 | | 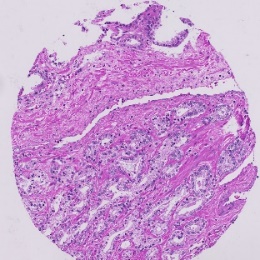 | H3 | 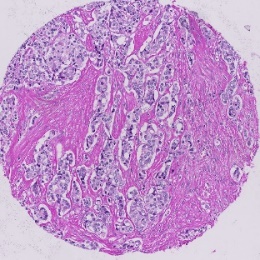 | 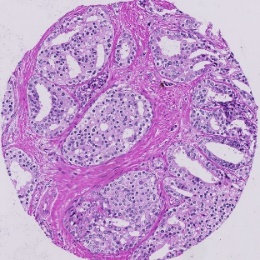 |
|  | R2 | 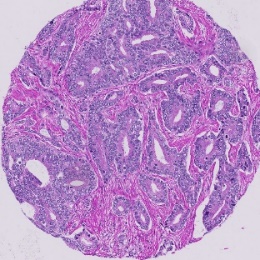 | | 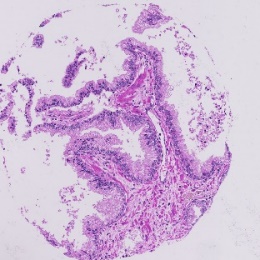 |  | 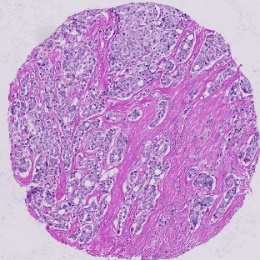 | 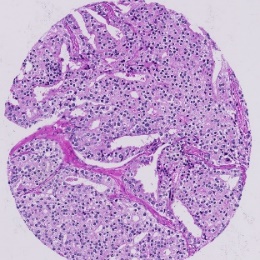 |

**Figure S10. Histology images of TA1 and paired TA2 in two replicates (R1 and R2) for selected patients. (a)** These three patients (H2, H4, M13) have highly correlated (paired) tumor areas at all molecular layers. **(b)** The two patients (M12, M14) have weakly correlated (paired) tumor areas at all molecular layers. **(c)** The two patients (M9, M17) have very similar (paired) tumor areas at the CNA and mRNA layers but not at the protein layer. **(d)** The two patients (M6, H3) have very similar (paired) tumor areas at the protein layer but not at the CNA and mRNA layers.

**Figure S11**

**a**

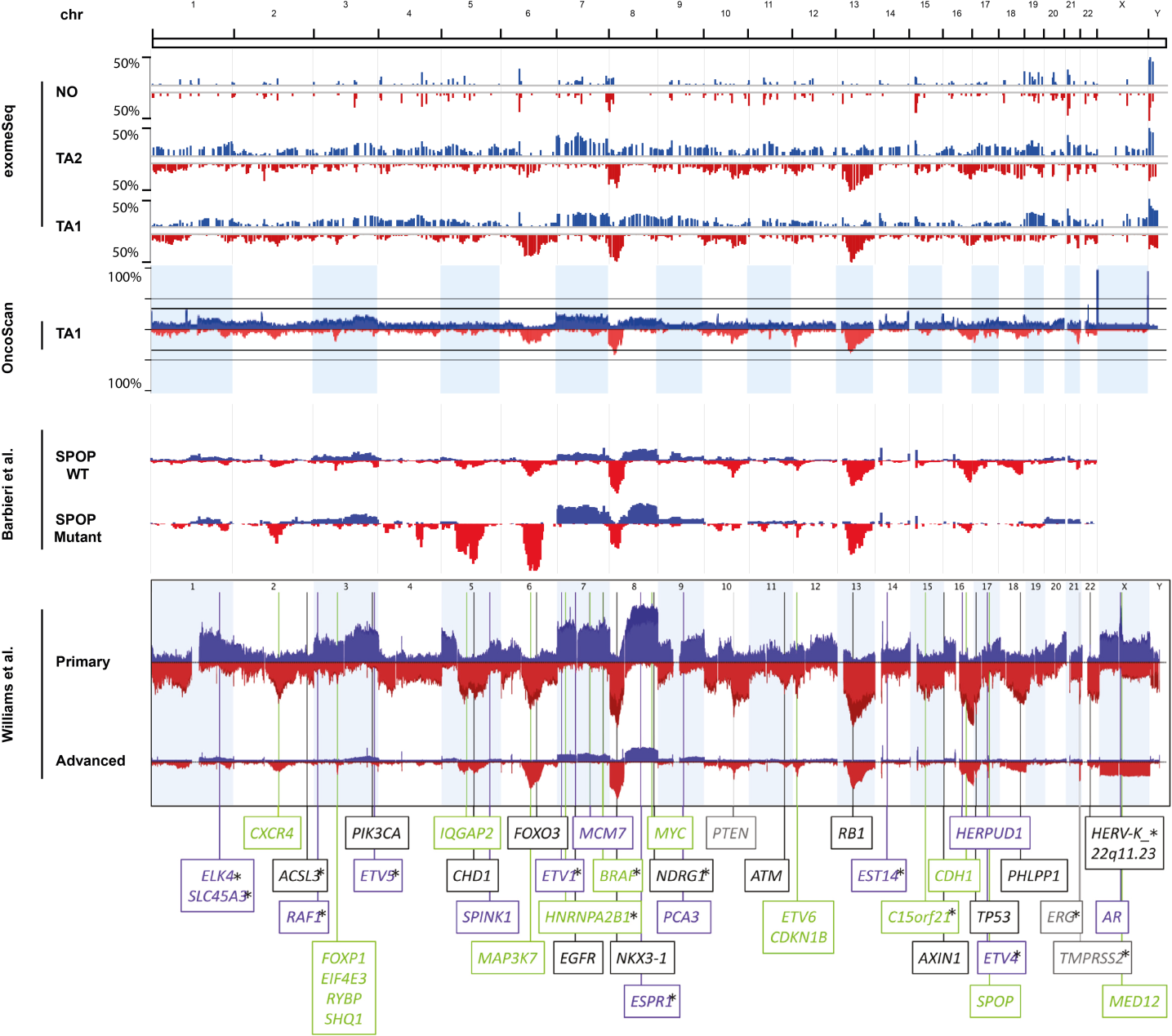

**Figure S11 (*continued*)**

**b**

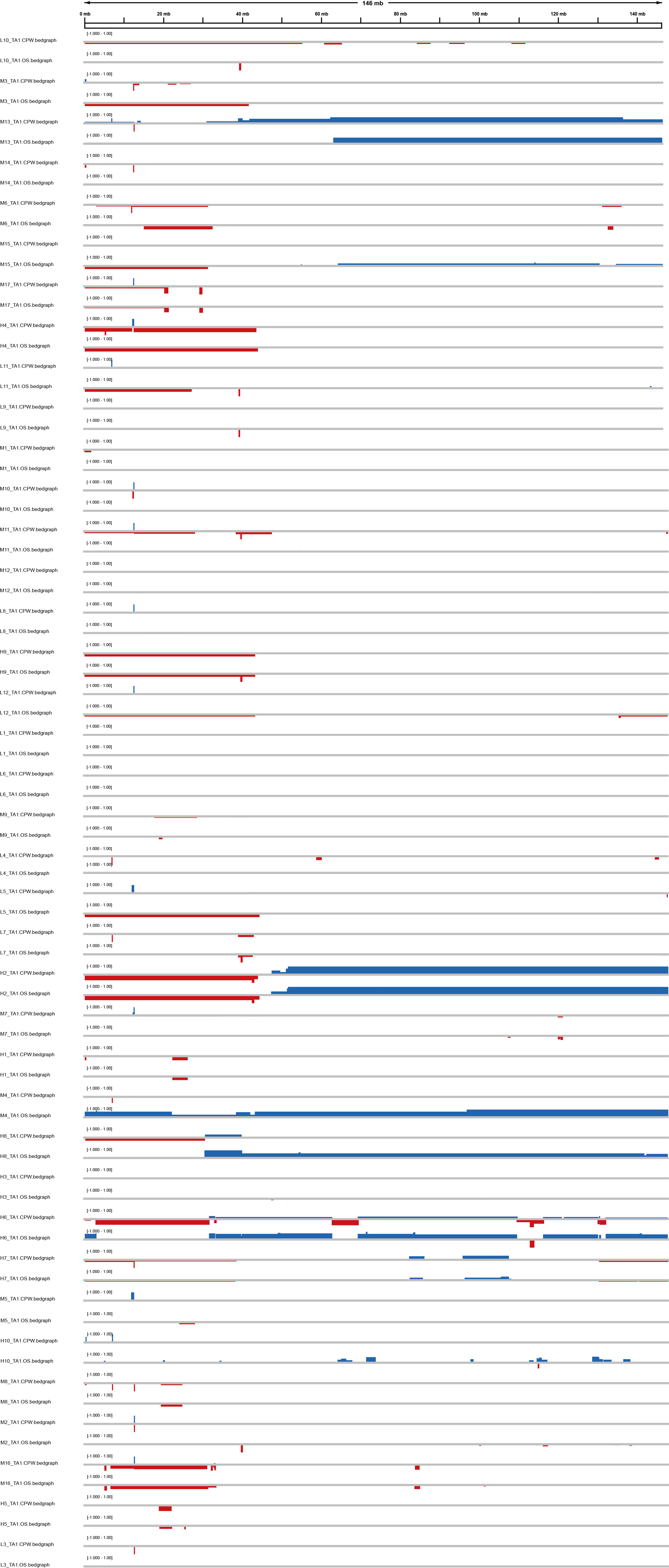

**Figure S11. Comparison of two methods for CNA-calling (exome-seq and Oncoscan). (a)** Recurrent CNAs in the PCa samples as detected in exome-seq data using the CopyWriteR software. Genome-wide frequencies for copy number gains (blue) and losses (red) are displayed for benign tissues (NO), TA, TA1 and secondary tumor areas (TA2). The lowest panel shows the genome-wide frequencies for copy number changes in all TA1 and TA samples as detected by the Affymetrix OncoScan platform showing that CNA results detected by sequencing- and array-based methods are comparable. To show that the resulting CNA frequency profiles are in agreement with previous studies, previously published CNA frequency profiles are displayed. Briefly, CNA frequency profiles for SPOP WT and mutant PCa samples are chosen from Barbieri et al. [1]. Finally, CNA frequency profiles are shown for primary and advanced PCa from the meta-analysis carried out by Williams et al [4]. In the latter panel the genomic positions of a list of PCa candidate driver genes are indicated. **(b)** Copy number gains (blue) and losses (red) in chromosome 8 of all TA1 and TA samples. CNAs were detected by exome-seq combined with CopyWriteR software and by Affymetrix OncoScan and displayed using Integrated Genome Viewer showing that application of both platforms yields fairly consistent results.

**Figure S12**

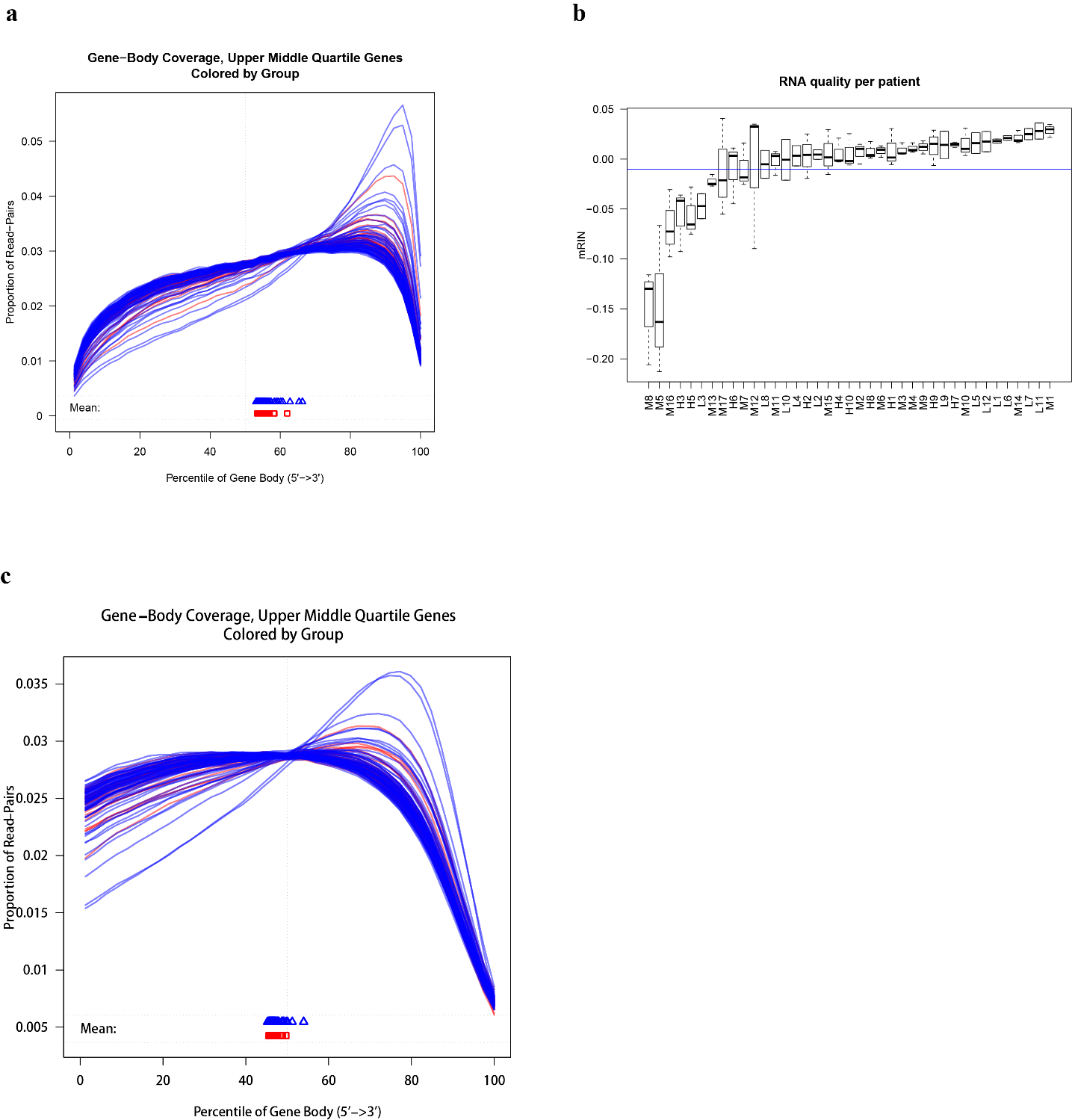

**Figure S12. Degradation of RNA samples.** **(a)** Gene body coverage of upper and middle quartile genes (by read count). Tumor samples are plotted in blue, benign samples are plotted in red. **(b)** mRIN scores per patient. Each box represents the mRIN score of all samples of the respective patient. The blue horizontal line separates highly degraded samples (mRIN < -0.01, *P* value < 0.1) from the rest. **(c)** Gene body coverage of upper and middle quartile genes after correction for 3’ bias. (Gene body coverage plots were generated using the R package QoRTs.)

**Figure S13**

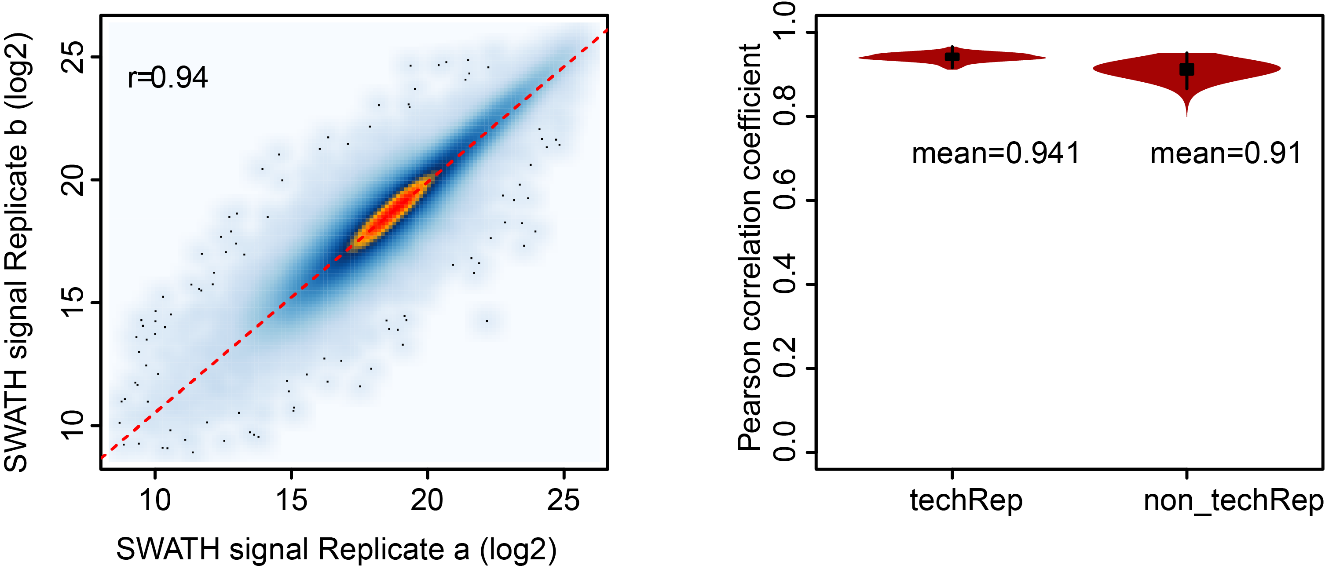

**Figure S13. Protein dataset quality analysis.** The technical reproducibility of duplicate SWATH analysis (left) and Pearson correlation coefficient of replicate samples and the non-replicate samples (right). Violin plots were made using R package vioplot.
